# Climate-driven fitness decline in Japanese Chum Salmon reshapes North Pacific salmon biogeography, with reproductive consequences: A Bayesian network analysis

**DOI:** 10.64898/2026.07.02.735760

**Authors:** Shuichi Kitada, Hirohisa Kishino

## Abstract

Japanese Chum Salmon (JCS) supported by one of the world’s largest hatchery programs have experienced severe declines in abundance, return rate, and egg size. To investigate the underlying mechanisms, we analyzed a 21-year time series of reproductive traits of age-4 hatchery fish from 13 rivers together with climate and salmon abundance data using a bootstrap-supported Bayesian network. Here, we assumed that environmental variables can affect the salmon populations, but not vice versa, and that there could be maternal effect on reproductive traits, but not the other way around. The final network identified a major pathway from NPGO3 through JCS abundance to Russian Chum Salmon (RCS) abundance, followed by pathways from RCS to female fork length (FL) and from FL to egg size. A separate maternal pathway linked FL to egg size through fecundity. All corresponding edges had edge strength >0.97, and standardized path coefficients were statistically significant. These findings suggest that climate-driven fitness decline in JCS has reshaped salmon biogeography across the North Pacific, with consequences for reproductive traits, including a decline in egg size.

## 1. Introduction

Understanding how climate change affects population dynamics is a central challenge in fisheries science (Cheung et al. 2021; Crozier et al. 2021; Cheung 2025). Fish are ectotherms, making their survival and physiological performance directly sensitive to water temperature. Anadromous Pacific salmon (*Oncorhynchus* spp.) have complex life histories (Hilborn et al. 2003; Crozier et al. 2008), exposing them to climatic variability operating across multiple spatial and temporal scales. Because they return to their natal rivers for spawning, they cannot avoid local warming conditions. In addition, they undertake extensive oceanic migrations across the North Pacific, exposing them to basin-scale environmental variability. Their semelparous life history further amplifies this sensitivity, as a single reproductive event determines lifetime fitness, making population productivity highly responsive to environmental conditions throughout the life cycle.

In many Pacific salmon species, long-term declines in abundance and reproductive traits have been reported (Lewis et al. 2015; Cline et al. 2019; Oke et al. 2020; Ohlberger et al. 2023; Malick et al. 2023; Kitada 2026), although conservation efforts may help to stabilize and increase salmon population abundance (Ford et al. 2025). These contrasting trends suggest that population responses to environmental change and management interventions are mediated through multiple biological mechanisms, including changes in life-history traits. However, the mechanisms linking environmental variability to biological responses remain poorly understood. This limitation arises in part because environmental drivers and biological processes are often examined separately, and most existing studies rely on association between explanatory and response variables rather than causal inference, obscuring how climatic effects propagate through complex life-history pathways. The challenge is further compounded by hatchery enhancement programs, which can influence the abundance, distribution, and life-history traits of salmon populations (Waples 1991; Naish et al. 2007; Cunningham et al. 2018; Riddell et al. 2025).

For more than a century, large-scale salmon hatchery programs have been implemented across the Pacific Rim (Naish et al. 2007; Kitada 2018), and hatchery-origin salmon have comprised approximately 40% of total adult and immature biomass in the North Pacific (Ruggerone and Irvine 2018). Although hatchery enhancement has increased fishery production (Hilborn and Eggers 2000; Kitada 2020), prolonged reliance on artificial propagation can induce eco-evolutionary feedbacks that may compromise life-history traits, reproductive success, genetic structure, and long-term sustainability (Waples and Drake 2004; Araki et al. 2007; Christie et al. 2012; Christie et al. 2014; Kitada and Kishino 2021; Malick et al. 2023; Kitada 2026).

Among life-history traits in salmonids, egg size plays a central role in shaping early survival, growth trajectories, and lifetime reproductive success, reflecting maternal investment strategies that balance fecundity, offspring quality, and environmental uncertainty (Beacham and Murray 1993; Einum and Fleming 1999, 2000; Fleming and Gross 1990; Heath et al. 1999, 2003). Consequently, variation in egg size is closely linked to fitness, population productivity, and adaptive capacity (Quinn et al. 1995; Chambers and Leggett 1996; Mousseau and Fox 1998). In a related study (Kitada 2026), synchronized declines in egg size, return rate (an index of post-release survival), and abundance in Japanese Chum Salmon (*O. keta*; hereafter, JCS) were demonstrated. Female fork length (FL) was positively correlated with egg size, and egg size was positively correlated with subsequent return rate. The estimated instantaneous exponential decay rate of JCS abundance was close to the relative reproductive success (RRS)-based estimate reported by Araki et al. (2007). Together, these findings suggest a long-term decline in fitness of JCS. However, the causal pathways linking environmental variability to changes in abundance, female body size, and egg size remain unresolved.

Here, we infer direct dependence relationships linking environmental variability, salmon abundance, female body size, and egg size using a Bayesian network framework. We analyzed a 21-year time series (1999–2019) of reproductive traits in JCS from 13 hatchery-enhanced rivers in Japan, previously analyzed in the related study (Kitada 2026), together with oceanographic and salmon catch data. We combined block bootstrap resampling with Bayesian network analysis to account for temporal dependence and synchronous variation among rivers. This framework allowed us to examine how environmental variability and salmon abundance were linked to changes in female body size and reproductive traits including egg size, with the North Pacific Gyre Oscillation (NPGO; Di Lorenzo et al. 2008) used as an index of basin-scale climate variability. By integrating long-term data within a hypothesized causal framework, this study provides an integrated view of how climate variability, salmon population dynamics, and reproductive traits are linked in overwhelmingly hatchery-supported JCS and the broader North Pacific.

## 2. Methods

### 2.1 Data

We analyzed global monthly mean sea surface temperature (SST) data at a spatial resolution of 0.25° × 0.25° for the period 1982–2022. Monthly mean SSTs for seasonal migration areas were obtained from the datasets monmean.goa.nc and monmean.ber.nc in Data S7 of Kitada et al. (2025), which were derived from the NOAA Optimum Interpolation (OI) SST V2 high-resolution dataset (sst.mnmean.nc; NOAA 2026). Monthly NPGO data for 1950–2025 were obtained from the Monthly Climate and Ocean Indices database maintained by the NOAA Physical Sciences Laboratory (NOAA 2026). Annual mean NPGO values were computed from the monthly data.

We used the trait data originally derived from the Fisheries Research and Education Agency (FRA) salmon database (FRA 2026). For the present study, we used fork length (FL), egg size, and fecundity of age-4 female Chum Salmon sampled from 13 hatchery-enhanced rivers in Japan during 1999–2019 (5 rivers in Hokkaido and 8 rivers in Honshu; Figs. S1, S2). Detailed descriptions of data collection and trait measurements are provided in the related study (Kitada 2026), which analyzed the same dataset. We also compiled Chum Salmon commercial catch and hatchery release statistics from the North Pacific Anadromous Fish Commission (NPAFC 2026a), spanning 1952–2025.

### 2.2 Ecological background for Bayesian network modeling

The ecological context of this study is based on the proposed migration routes and seasonal distribution areas of JCS by Kitada et al. (2025) through a synthesis of previous studies (Fig. 1). Fig. 1 shows that JCS returning to Japan at age *i* are assumed to have experienced environmental conditions in the Bering Sea during *i* summers and in the Gulf of Alaska during (*i* − 1) winters during their marine life history. Few JCS matured and returned at age 2, although the proportion of age-3 fish has increased slightly in recent years. Age-4 fish remained the dominant age class throughout the study period, accounting for 40–77% of returning fish, followed by age-5 fish (15–39%). Fish aged 6 years or older were rare (see Fig. 5 in Kitada 2026).

**Fig. 1.**
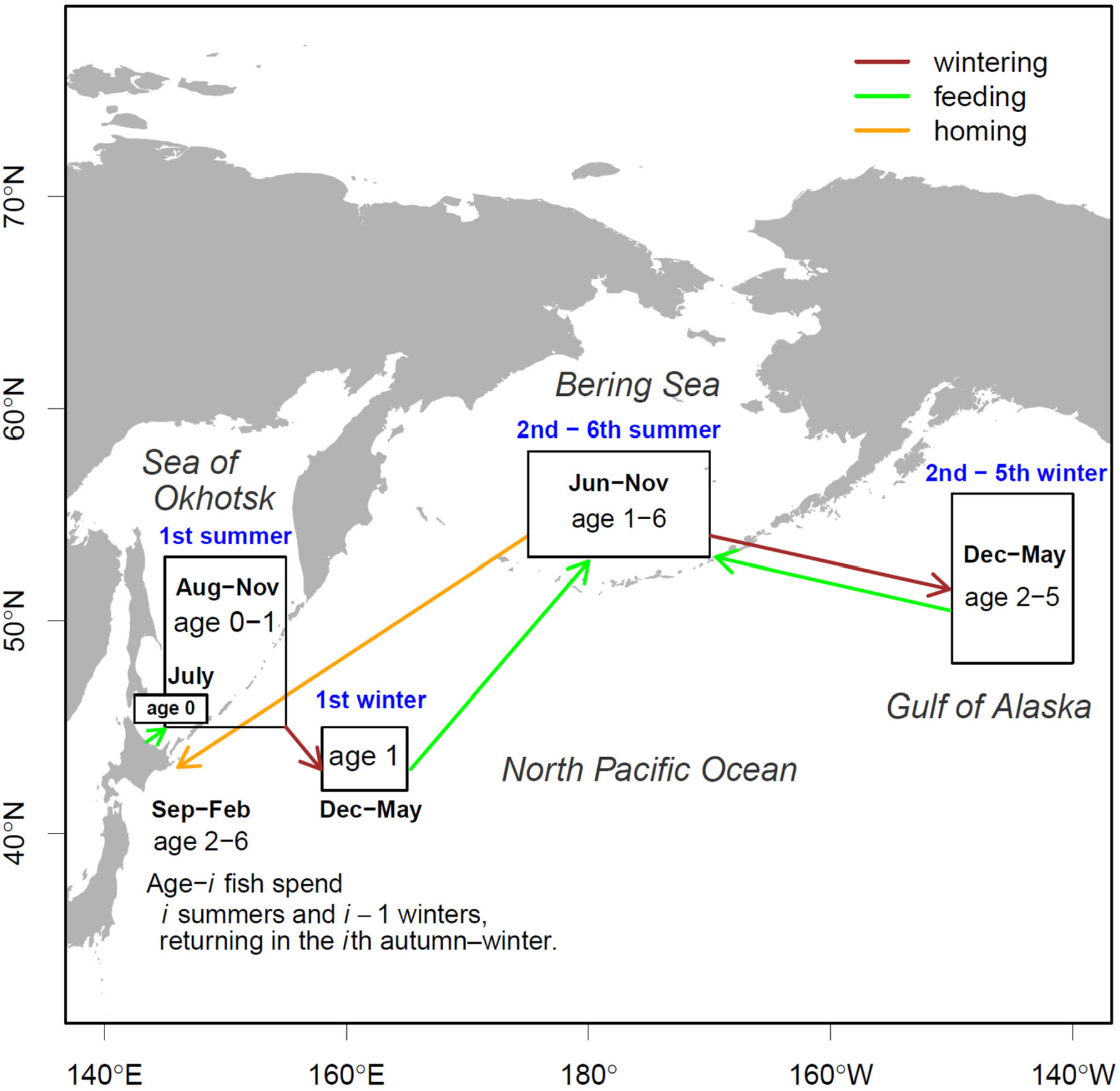
Ecological background for the Bayesian network analysis. Proposed migration routes and seasonal distribution areas of Japanese Chum Salmon based on previous studies. Adapted from Fig. 1b in Kitada et al. (2025).

We focused on age-4 females for the analysis of reproductive characteristics for several reasons. The trait dataset comprised six age groups: age-2 (*n* = 17), age-3 (*n* = 1634), age-4 (*n* = 12428), age-5 (*n* = 8864), age-6 (*n* = 619), and age-7 (*n* = 8). Age-4 fish were substantially more abundant than fish in the other age groups. Because Chum Salmon returning to Japan at age *i* are assumed to have experienced environmental conditions over multiple years of their marine life history (Fig. 1), the environmental history becomes increasingly complex with increasing age. Age 4 thus provided a balance between a large sample size and a relatively simple marine environmental history. In addition, the temporal trends in female FL and egg size were similar between age-4 and age-5 fish (Kitada 2026), supporting the use of age-4 fish as a representative age class for examining long-term changes in reproductive characteristics. Restricting the analysis to a single age class also minimized potential confounding effects arising from age-related variation in reproductive traits.

The North Pacific Gyre Oscillation (NPGO) represents variability in large-scale ocean circulation and associated physical oceanographic processes, including wind-driven upwelling and horizontal transport (Di Lorenzo et al. 2008). These processes can influence nutrient availability, planktonic ecosystem dynamics, and marine productivity, with NPGO variability having been associated with productivity of Pacific salmon stocks (Malick et al. 2017; Davis et al. 2023), density-dependent competition (Debertin et al. 2017), and body-size decline in Pacific salmon (Jeffrey et al. 2017; Oke et al. 2020). We therefore considered the NPGO a biologically relevant indicator of climate-driven environmental variability potentially affecting salmon population dynamics across the North Pacific.

Salmon mortality can be highest during the first year at sea (Beamish and Mahnken 2001). A previous study suggested that substantial mortality of hatchery-released fry occurs in Japanese coastal waters (Kitada et al. 2025). Accordingly, we hypothesized that variation in JCS abundance is strongly influenced by environmental conditions experienced during the first year of marine life. Most Chum Salmon returning to Japan are hatchery-released fish or wild-born descendants of hatchery-origin fish (Kitada 2000, see Section 4.4). Therefore, returning adults were linked to environmental conditions in their release year, representing conditions experienced during their first year at sea.

JCS share major oceanic habitats and seasonally overlap in feeding grounds with RCS (Urawa et al. 2022). Previous studies reported negative associations between RCS catches and JCS catches (Kitada 2018). Another study reported that return rates of JCS released from the Hokkaido Sea of Japan region were negatively associated with RCS catch (Kitada et al. 2025). Together, these studies suggest that environmental variability and interactions with RCS jointly influence the population dynamics of JCS.

Based on the ecological background described above, we hypothesized that the seven variables included in the Bayesian network were linked through biologically plausible pathways involving climate-driven changes in salmon abundance, interspecific interactions, geographic variation in river-mouth latitude, and associated changes in female body size and reproductive traits, including egg size. Definitions, abbreviations, units, and data sources for all variables are summarized in Table 1. The following sections describe the rationale for selecting these variables and the biological and ecological knowledge underlying their inclusion in the Bayesian network, covering environmental variables and climate indices (Section 2.3), abundance indices and hatchery releases (Section 2.4), and reproductive traits and geographical variation (Section 2.5).

**Table 1.** Summary of covariates used in the Bayesian network analyses, including their categories, abbreviations, definitions, units, and sources. *t* denotes the return year of age-4 hatchery Chum Salmon (1999–2019).

| Category | Covariate | Abbrev. | Description | Unit | Data source |
| --- | --- | --- | --- | --- | --- |
| River environment | River latitude | River | River latitude of sampling site | °N | Kitada (2026) |
| Marine environment (physical) | North Pacific Gyre Oscillation (NPGO) | NPGO3 | NPGO in the release year of age-4 hatchery fish ($t-3$) returned in year $t$. | index | NOAA (2026) |
| Marine environment (biological) | Japanese Chum Salmon abundance | JpChumN | Total return of Japanese Chum Salmon in year $t$. | indiv. | NPAFC (2026a) |
| | Russian Chum Salmon abundance | RusChumN | Total return of Russian Chum Salmon in year $t$. | indiv. | |
| Reproductive traits | Fork length | FL | Mean fork length of age-4 female Japanese Chum Salmon returned in year $t$. | cm | Kitada (2026) |
| | Fecundity | Fecund | Number of eggs per age-4 female Japanese Chum Salmon returned in year $t$. | eggs | This study |
| | Egg size | EggSize | Mean egg weight of age-4 female Chum Salmon returned in year $t$. | g | Kitada (2026) |

### 2.3. Environmental variables and climate indices

Because the NPGO is defined at the basin scale, it cannot be calculated separately for the marine distribution ranges of JCS. We therefore used the NPGO index as an indicator of broad-scale oceanographic conditions. To examine the relationship between NPGO and broad-scale thermal variability, we calculated mean SSTs for 16 geographic areas (Fig. 2A) covering the principal marine distribution of Chum Salmon (Dunmall et al. 2022; Langan et al. 2024). Monthly mean SSTs for each area were calculated from the NOAA OI SST V2 dataset (sst.mnmean.nc) for 1982–2022 using the `fldmean()` function in the `cdo-2.1.1` package (Max Planck Institute 2023). Annual unweighted mean SST remained relatively stable until around 2010 in the 16 areas, but increased thereafter (Fig. S3). Because the areas differed substantially, annual mean SST across the 16 areas was calculated using geographic area (Table S1) as a weight. Thus, for each year, the area-weighted mean SST was calculated as the sum of the mean SST of each area multiplied by its geographic area, divided by the total area of the 16 areas. We also calculated annual mean SST assigning equal weight to each region. The area-weighted and unweighted mean SST indices were closely correlated (*r* = 0.98, 1991–2022) (Fig. S4).

**Fig. 2.**
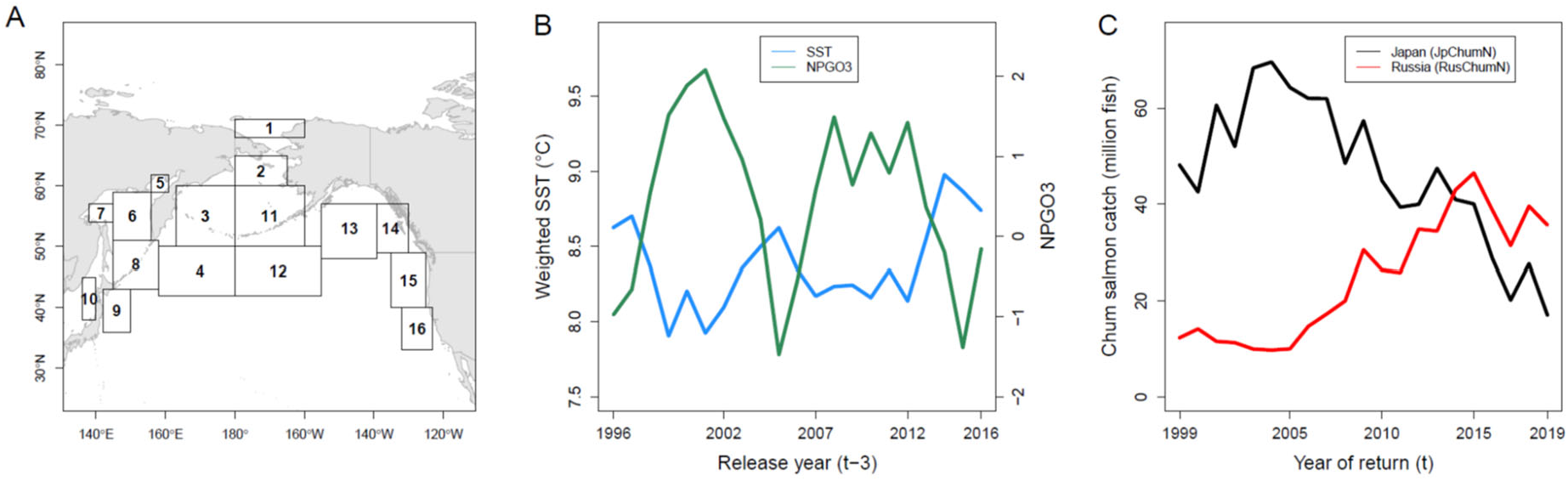
Physical and biological environments in the major Chum Salmon distribution areas. (A) Map showing the 16 areas used to calculate annual mean sea surface temperature (SST) (see Table S1). Panels B and C show time series during the Bayesian network analysis period (1999– 2019). (B) North Pacific Gyre Oscillation (NPGO3) and weighted annual mean SST across the 16 areas in the release year of age-4 fish (*t*−3). (C) Japanese and Russian Chum Salmon catch in year *t* (*r* = −0.72, *t* = −4.59, df = 19, *p* < 0.001). Here, *t* denotes the return year of age-4 Chum Salmon and *t*−3 denotes the corresponding release year.

Because our Bayesian network analysis related reproductive characteristics and number of age-4 fish returning during 1999–2019 to environmental conditions during their release year, NPGO was lagged by three years, corresponding to the interval between hatchery release (*t* − 3) and return of age-4 fish in year *t*. We denote it as NPGO3. The suffix “3” denotes this 3-year lag. We examined the relationship between NPGO3 and the weighted annual mean SST across the 16 areas during the release years (1996–2016) corresponding to age-4 returns in the Bayesian network study period (1999–2019) (Fig. 2B). NPGO3 was strongly negatively correlated with the weighted annual mean SST (*r* = −0.83; Fig. S5). We therefore used NPGO3 as a basin-scale index of broad-scale oceanographic conditions during the first year at sea. Because NPGO3 and SST provided largely overlapping information, SST was not included as a separate variable in the Bayesian network.

### 2.4. Abundance indices and hatchery release

We used annual commercial catch (NPAFC 2026a) as an index of returning salmon abundance. Most JCS catch is taken by coastal set-net fisheries (MAFF 2026a), indicating that commercial catch largely reflects the harvest of returning salmon in coastal waters. The high-seas salmon fishery has been closed since 1992 (Morita et al. 2006). Thus, JCS fisheries have relied largely on the capture of returning salmon in coastal waters. According to the national fisheries census conducted every five years (MAFF 2026b), the number of fishing enterprises engaged in salmon set-net fisheries averaged 606 during 2003–2023. This number increased through 2013 and remained above the 2003 level in subsequent census years (Fig. S6). The continued decline in JCS catch since its peak in 2004 (Fig. 2C), despite no decline in fishing effort as indicated by the number of set-net fishing enterprises, strongly suggests a decline in the abundance of returning JCS. Russian Chum Salmon (hereafter; RCS) commercial catch data were also obtained from NPAFC. In Russia, salmon catches are regulated according to the abundance of returning salmon (NPAFC 2026a). These characteristics support the use of commercial catch as an index of annual salmon abundance in both countries.

Hatchery-related factors, including the number, size, and quality of released fry, can affect salmon abundance. In Japan, the number of hatchery-produced fry released peaked in the 1980s and declined slightly thereafter following the termination of ineffective stocking programs (Miyakoshi et al. 2013), but remained above 1.5 billion per year during 1996–2016, the period corresponding to our study period (1999–2019) with a three-year lag (Fig. 2 in Kitada et al. 2025). Despite the sustained high level of releases, JCS catch declined substantially after 2005. This decline coincided with declines in egg size and return rate (Fig. 6 in Kitada 2026). In Russia, total Chum Salmon catch increased sharply after around 2005 despite a gradual increase in hatchery releases, but declined after 2015 as hatchery releases increased markedly (Fig. S7). Temporal relationships between hatchery releases and catch also varied among Russian regions. Nevertheless, catch declines after around 2015 were observed across all major production regions despite continued releases. Notably, in the Kuriles and Eastern Sakhalin, which had the largest release programs and experienced sharp increases in releases from 2020 onward, recent catch trends diverged markedly: catch declined sharply in the Kuriles but increased sharply in Eastern Sakhalin. These contrasting patterns indicate that variation in catch, and by implication returning salmon abundance, cannot be explained by the number of hatchery releases alone.

Given these patterns, we included population abundance metrics as biological proxies reflecting the combined effects of hatchery production, marine ecosystem conditions, and potential density-dependent interactions. JCS abundance was represented by annual catch in number (JpChumN). Annual catches (in number) of RCS (RusChumN) were included to account for potential ecological interactions between sympatric Chum Salmon populations (Fig. 2C). After peaking at nearly 70 million fish in 2004, JCS catches (JpChumN) declined continuously and have remained at low levels. In contrast, RCS catches (RusChumN) remained relatively low, at approximately 10 million fish annually, until 2005, but then increased steadily through 2015, reaching a level comparable to JCS catches. This temporal pattern contrasted with the decline observed in Japan. Russian catches subsequently also declined. By 2015, RCS catches had exceeded JCS catches, and the difference between them widened thereafter. JpChumN was strongly negatively correlated with RusChumN during the study period (*r* = −0.72).

### 2.5. Reproductive traits and geographical variation

Reproductive trait data consisted of age-4 female JCS sampled from 13 hatchery-enhanced rivers in Japan during 1999–2019 (*n* = 12428), after outliers excluded (Kitada 2026). Female fork length (FL) and fecundity (Fecund) were included in the Bayesian network as maternal reproductive traits potentially influencing egg size (EggSize). The latitude of the natal river mouth (River) was included as an environmental covariate to account for the latitudinal variation in female FL, egg size and fecundity (Fig. S2).

Pairwise scatter plots of individual fork length (FL), egg size (EggSize), and fecundity (Fecund) for age-4 female Chum Salmon pooled across the 13 rivers showed moderate but statistically significant correlations among the three traits. FL was positively correlated with EggSize (*r* = 0.31) and Fecund (*r* = 0.56), whereas EggSize was negatively correlated with Fecund (*r* = −0.20) (Fig. 3). To examine whether these relationships were consistent across years, we compared scatterplots for 1999, 2009, and 2019 in two rivers with relatively large sample sizes: the Tokushibetsu River (*n* = 892) and the Nishibetsu River (*n* = 1071) (Figs. S8, S9). The patterns in these two rivers were broadly consistent with those observed in the pooled dataset (Fig. 3). In both rivers, EggSize declined over the study period, whereas the decline in FL was less pronounced and no substantial decline in Fecund was apparent. These patterns indicate that the temporal decline in EggSize was not accompanied by comparable declines in female body size or fecundity.

**Fig. 3.**
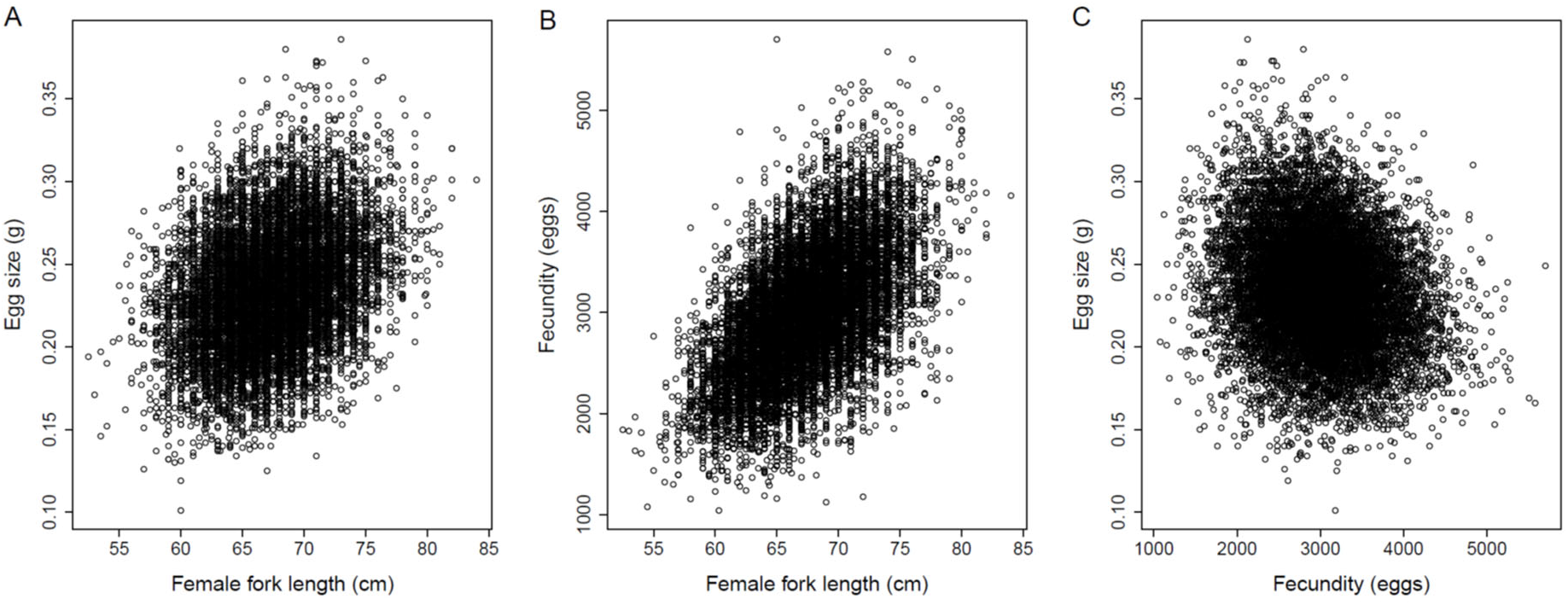
Pairwise scatter plots of fork length (FL), egg size, and fecundity of age-4 female Chum Salmon from 13 hatchery-enhanced rivers (see Fig. S1). Plots are based on the screened dataset used for the Bayesian network analysis after excluding apparent outliers (*n* = 12,428). (A) FL was positively correlated with egg size (*r* = 0.31, *t* = 36.3) and (B) fecundity (*r* = 0.56, *t* = 74.5), whereas egg size was negatively correlated with fecundity (C) (*r* = −0.20, *t* = −22.7) (all tests: df = 12,426, *p* < 0.0001).

### 2.6 Bayesian network inference and edge selection

A Bayesian network is a probabilistic graphical model represented as a directed acyclic graph (DAG), in which variables are represented as nodes and conditional dependencies among variables are represented as directed edges (arcs) (Pearl 1985; Scutari 2010; Scutari and Denis 2021). In this study, the variables listed in Table 1 were represented as nodes, and directed edges represented conditional relationships between pairs of nodes. Rather than testing a prespecified set of relationships among variables, our objective was to infer the network structure and identify pathways consistent with the conditional dependencies in the data, subject to constraints based on prior biological and ecological knowledge. Bayesian network analysis involves two main steps: network structure learning and parameter estimation. We used the hill-climbing algorithm implemented in the `bnlearn` package in R (Scutari 2010), which uses the Bayesian information criterion (BIC) as the scoring function, to learn the network structure from the data. Biological constraints were incorporated by specifying a blacklist of prohibited directed edges based on a priori biological and ecological knowledge (Sections 2.2–2.5). These constraints were the only a priori restrictions imposed on the network structure; all remaining relationships were inferred from the data. Given the learned network structure, the parameters of the network were estimated. We used bootstrap resampling to evaluate the robustness of the inferred network structure and the uncertainty in parameter estimates. Structural robustness was assessed from the frequency with which edges were selected across bootstrap samples, whereas parameter uncertainty was evaluated from the distribution of estimated coefficients across bootstrap samples. A brief glossary of Bayesian network terms used in this study is provided in the Supplementary Information to facilitate interpretation of the results.

A directed path is defined as a sequence of directed edges linking two nodes through one or more intermediate nodes. We used a Gaussian Bayesian network, in which the conditional linear relationships between continuous variables are modeled using Gaussian distributions. Because all variables were standardized before model fitting, the regression coefficients associated with the directed edges represent standardized path coefficients (β), which quantify the direction and magnitude of the fitted directional relationships. Unlike conventional structural equation modeling (SEM), in which hypothesized relationships are typically specified in advance, Bayesian network structure learning allows the network structure to be inferred from the data while incorporating biologically reasonable constraints.

We analyzed seven variables (nodes), including physical environmental variables (NPGO3 and River), population abundance metrics (JpChumN and RusChumN), and maternal traits (FL, Fecund, and EggSize) (Table 1). Directional constraints were imposed to exclude biologically implausible relationships. Environmental variables were assumed not to be influenced by salmon abundance or maternal traits, and salmon abundance was assumed not to be influenced by maternal traits. Because reproductive traits were measured on individual females, egg size and fecundity were not permitted to influence female fork length (FL), as egg size in salmonids is determined by maternal characteristics (Beacham and Murray 1993; Fleming and Gross 1990). These assumptions were implemented as biological constraints using a blacklist (Table S2).

Although our analyses are based on individual-level data, annual mean female FL and egg size show synchronous variation among rivers (Kitada 2026), implying that individual observations are not fully independent and that the effective sample size can be smaller than the total number of observations. To account for temporal dependency among rivers, we performed 1000 block bootstrap resamplings in which entire years were resampled, retaining all individuals within each sampled year. Bootstrap support for each edge was assessed across 1000 block-bootstrap replicates. For each pair of nodes, the bootstrap support probabilities for the inferred direction (boot.p) and the opposite direction (opp.boot.p) were calculated. Edge strength was defined as boot.p + opp.boot.p, representing bootstrap support for the presence of an edge irrespective of its direction. Direction probability was defined as boot.p / (boot.p + opp.boot.p), representing the proportion of bootstrap support for the edge that favored the inferred direction.

To obtain an initial reference for selecting an edge-strength threshold, we first estimated edge strengths using the standard bootstrap procedure implemented in the `boot.strength()` function in the `bnlearn` package. This preliminary analysis used the same hill-climbing algorithm, blacklist constraints, and number of bootstrap replicates (1000) as the main analysis, but did not account for temporal dependence among observations from different rivers. We then used the `inclusion.threshold()` function to obtain an initial inclusion threshold. For the main analysis, which accounted for temporal dependence, we examined the resulting distribution of edge strengths and selected a threshold of 0.6 because the distribution did not show a clear separation between strongly and weakly supported edges (see Section 3.1). We also required a direction probability >0.5 as a majority-rule criterion, indicating that the inferred direction was recovered more frequently than the opposite direction across bootstrap replicates (Felsenstein 1985).

### 2.7 Final Bayesian network

For each directed edge retained under the specified thresholds, we calculated the mean coefficient and its 95% confidence interval using only bootstrap networks in which that directed edge was present (the nonzero-mean approach). This provides a conditional estimate of the coefficient given that the edge is present, rather than a marginal estimate that incorporates uncertainty in edge inclusion. To assess the effect of this choice, we also calculated the marginal mean and its 95% confidence interval across all bootstrap networks, assigning a coefficient of zero when the directed edge was absent.

The final network was visualized using the `igraph` package in R (Csardi and Nepusz 2006). Standardized path coefficients (β) for the retained edges were summarized using the mean and standard deviation (SD) of the non-zero estimates from the first-stage bootstrap (the nonzero-mean approach). Their 95% confidence intervals were calculated using the normal approximation (mean ± 1.96 SD), based on these non-zero estimates. The final network was visualized using the mean non-zero standardized path coefficients.

### 2.8 Mediated effects along selected pathways

To quantify direct and indirect (mediated) effects along selected pathways, we used the final Bayesian network structure obtained from the first-stage bootstrap as the fixed network structure and performed a second-stage bootstrap with 1000 replicates. Years were resampled as blocks, with all individuals within each sampled year retained, thereby preserving temporal dependence and synchronous variation among rivers. In each replicate, standardized path coefficients and mediated effects were estimated conditional on the fixed network structure. Thus, the estimated effects were conditional on the final network structure.

Standardized path coefficients for the 14 retained edges were estimated in each bootstrap replicate. The standardized path coefficient of each direct edge was taken as the corresponding direct effect. For selected indirect pathways, indirect effects were calculated as the product of the standardized coefficients along the constituent edges (e.g., the product of the FL → Fecund and Fecund → EggSize coefficients). The bootstrap distributions of these products were used to estimate their means and 95% confidence intervals.

All statistical analyses were performed in R (R Core Team 2026).

## 3. Results

### 3.1 Bayesian network inference and edge selection

Block bootstrap resampling across years (1000 replicates; *n* = 12428 individuals per replicate) was used to estimate the strengths and direction probabilities of edges among seven nodes. Edge strengths were calculated for all 21 possible pairs of nodes [7 × (7 − 1)/2]. Direction probabilities were estimated for the 23 directed edges recovered in at least one bootstrap replicate. The remaining 19 of the 42 possible directed edges were never recovered and therefore had bootstrap support of zero. No identical network structure was obtained across 1000 bootstrap replicates.

To determine an appropriate edge-inclusion threshold, we first performed a preliminary bootstrap analysis without block resampling. This analysis yielded an inclusion threshold of 0.314. The distribution of edge strengths across all 21 possible variable pairs showed a clear gap above this value, with edge strengths ≥0.9 occurring above the gap (Fig. S10). In contrast, when years were resampled as blocks to account for temporal dependence among rivers, edge strengths showed a more gradual decline (Fig. 4A). Of the 21 possible edges, 14 had edge strengths ≥0.665 (Table S3), whereas the remaining seven had edge strengths ≤0.494. We therefore adopted a conservative threshold of 0.6, which separated these two groups, and retained 14 edges with edge strength >0.6 and direction probability >0.5 in the final Bayesian network.

**Fig. 4.**
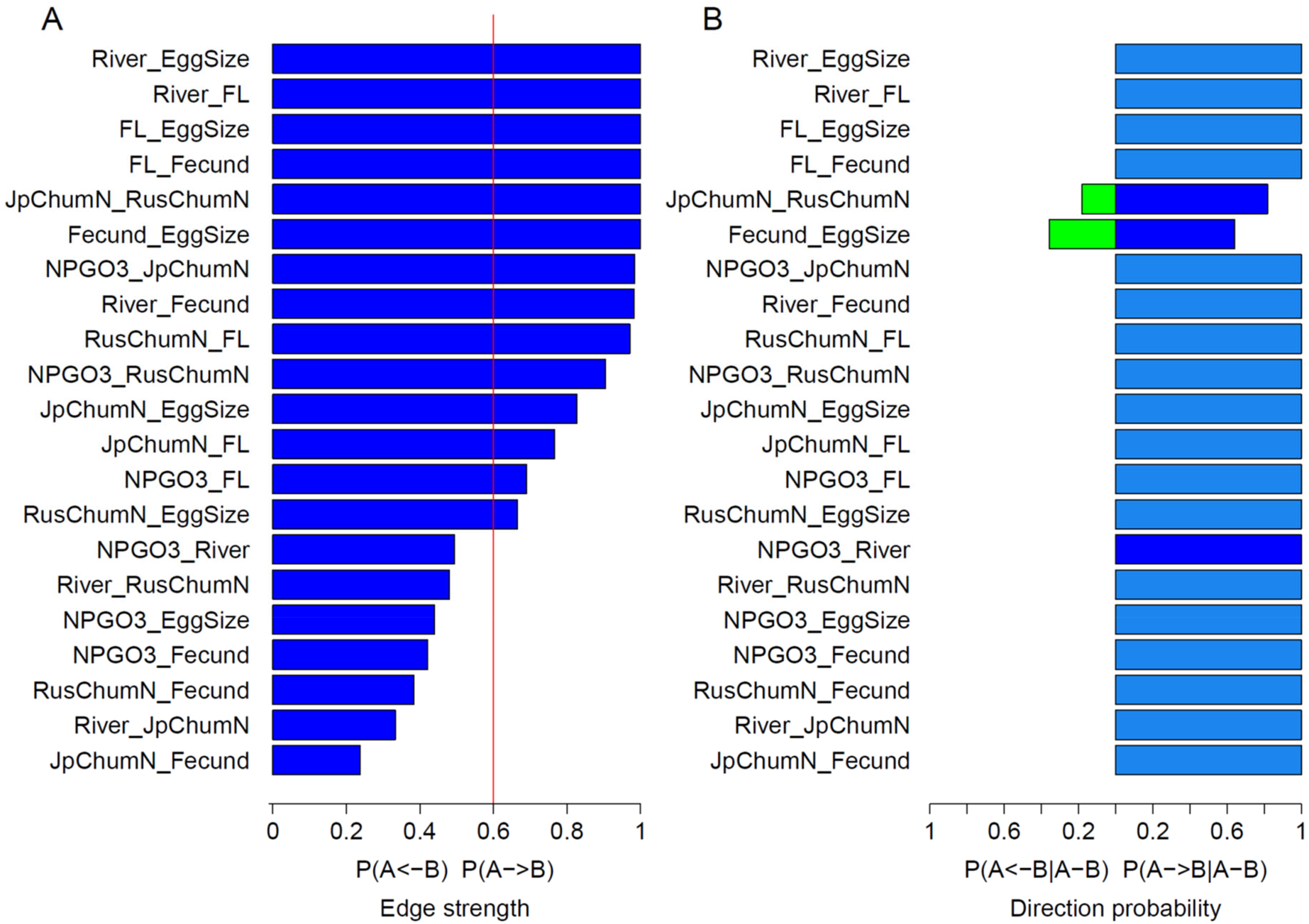
Bootstrap support for Bayesian network edges estimated from 1,000 block-bootstrap replicates, in which years were resampled while retaining all individuals (n = 12,428) within each year, thereby accounting for temporal dependence and synchronous variation among rivers. (A) Edge strength. The vertical red line indicates the edge-strength threshold (0.6). Edges with edge strength > 0.6 and direction probability > 0.5 were retained for construction of the final Bayesian network (see Fig. 5). (B) Edge direction probability. Blue and green bars represent edge direction probabilities estimated without biological constraints, whereas light blue bars represent the corresponding estimates obtained under the biological constraints defined in Table S2. Green bars indicate edges with direction probabilities < 0.5 and were therefore excluded from the final network. Biologically constrained estimates (light blue) were used only for edges that also satisfied the edge-strength criterion (> 0.6).

**Fig. 5.**
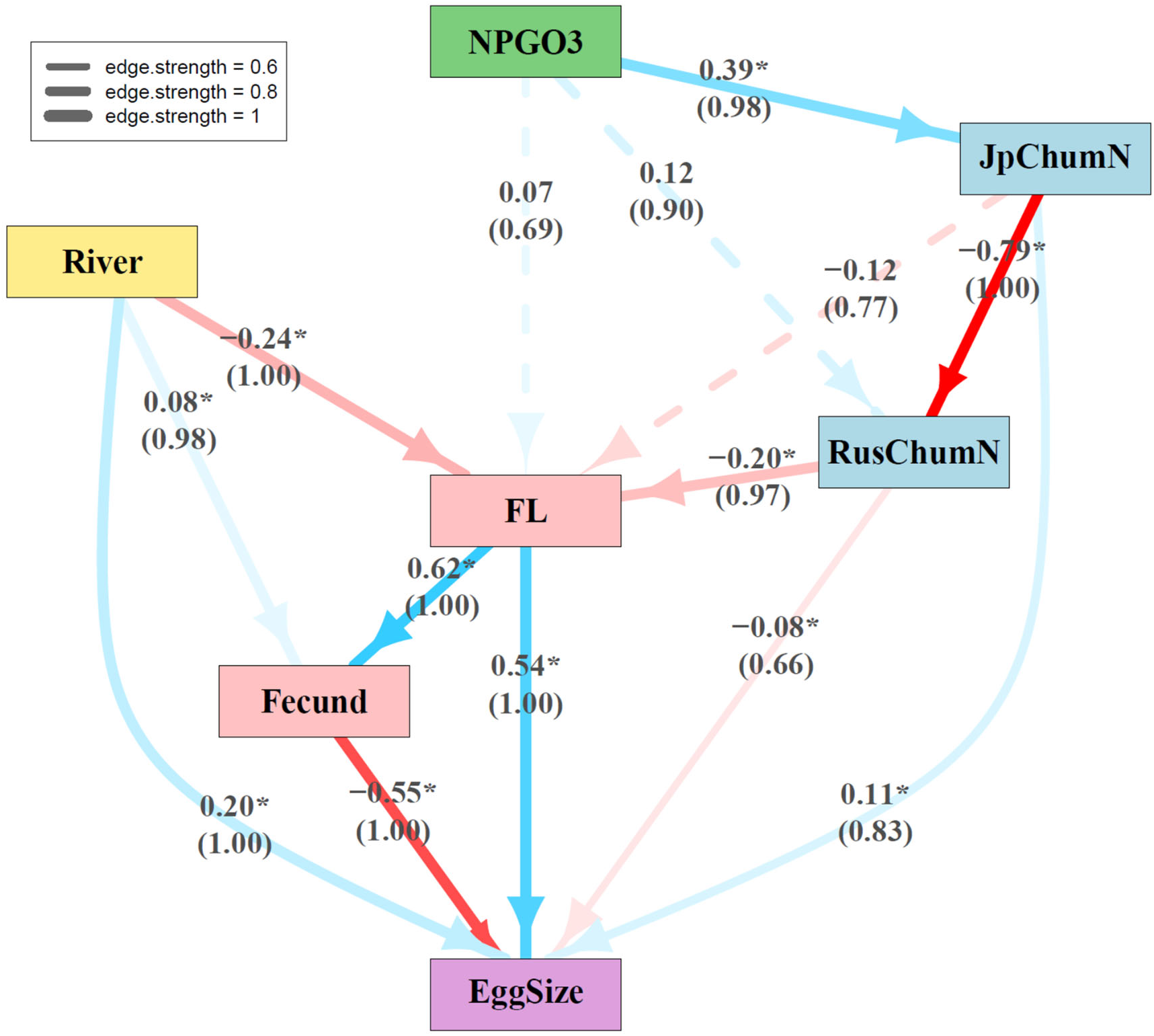
Final bootstrap-supported Bayesian network inferred from 1,000 block bootstrap replicates in which years (1999–2019) were resampled while retaining all individuals within each year (*n* = 12,428), thereby accounting for temporal dependence. Edges with edge strength > 0.6 and direction probability > 0.5 were retained for construction of the final Bayesian network, resulting in a network with seven nodes and 14 directed edges (see Fig. 4). The width of each edge is proportional to edge strength, and the size of the arrowhead is proportional to direction probability. Numbers shown above the edges are standardized path coefficients, and numbers in parentheses below indicate edge strengths. Asterisks indicate edges whose bootstrap 95% confidence intervals for standardized path coefficients excluded zero (see Fig. 6). Dashed lines indicate edges whose bootstrap 95% confidence intervals for standardized path coefficients included zero. Color intensity is proportional to absolute standardized path coefficient.

**Fig. 6.**
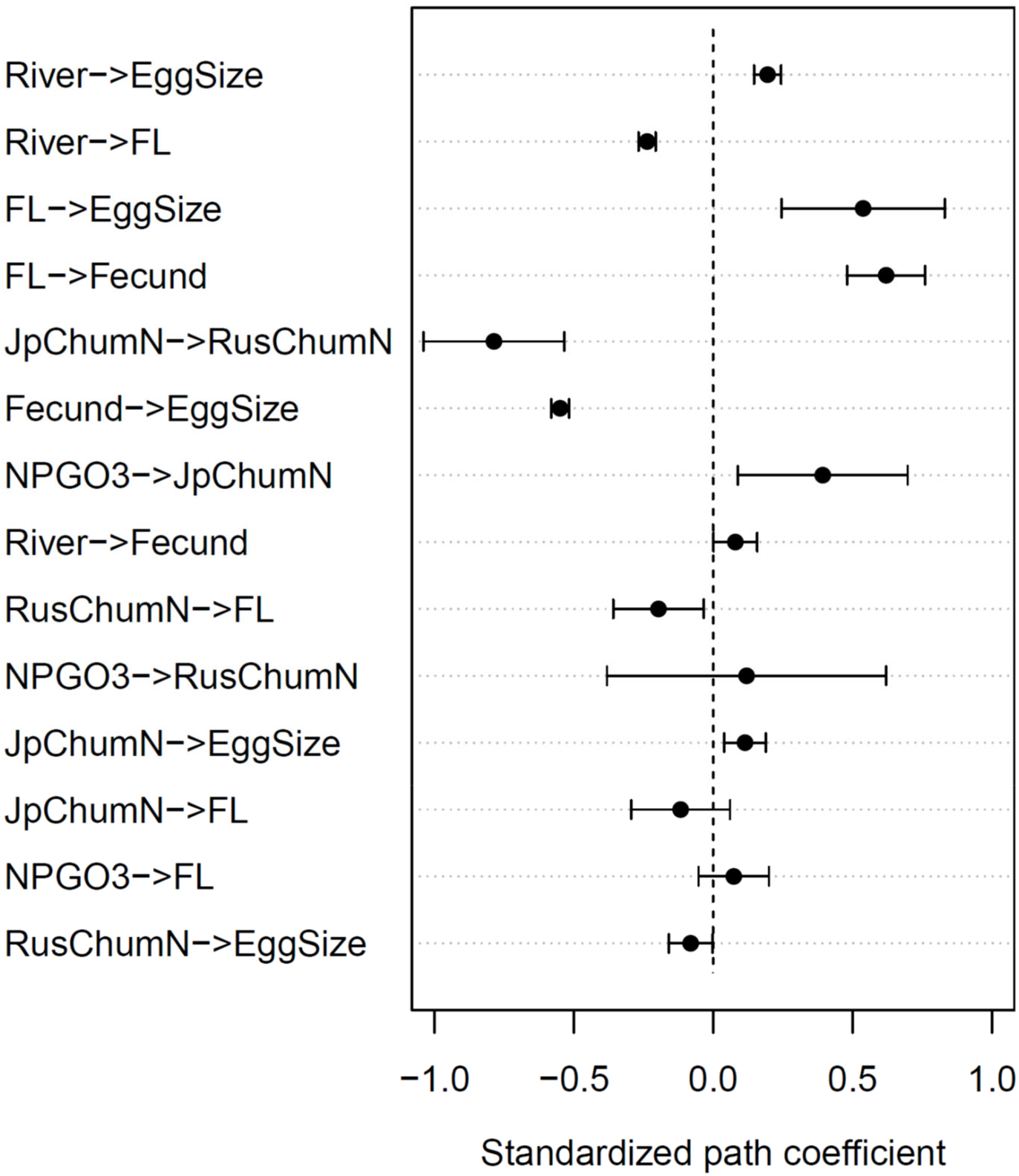
Bootstrap-averaged nonzero-mean standardized path coefficients (β) for 14 directional edges in the final Bayesian network (see Fig. 5). Dots indicate bootstrap means, and horizontal bars represent 95% bootstrap confidence intervals based on the normal approximation (mean ± 1.96 SD) across 1,000 block bootstrap replicates in which years (1999–2019) were resampled while retaining all individuals within each year (*n* = 12,428). Edges whose confidence intervals exclude zero were considered statistically supported.

Among the 23 directed edges recovered by bootstrap resampling, 14 (shown in blue and light blue) met both criteria of edge strength >0.6 and direction probability >0.5 (Fig. 4). Direction probabilities were 1.00 for 12 of the 14 retained edges (light blue in Fig. 4B), reflecting the biological constraints imposed by the blacklist (Table S2). The remaining two edges, JpChumN

→ RusChumN (direct.p = 0.82) and Fecund→ EggSize (direct.p = 0.64), were inferred without directional constraints and are shown in blue.

The marginal mean and nonzero-mean standardized path coefficient estimates were highly correlated (*r* = 0.94), indicating that the two approaches produced broadly similar estimates (Fig. S11). For 19 of the 23 directed edges, for which the direction probability was 1.0, the two point estimates and their 95% confidence intervals were nearly identical. Differences were observed for the two directed edges with lower direction probabilities, JpChumN → RusChumN and Fecund → EggSize (green dots in Fig. S11). The remaining two directed edges, RusChumN → JpChumN and EggSize → Fecund, had direction probabilities below 0.5 and were therefore excluded from the final network. Because our objective was to estimate the strength of the relationships represented by the directed edges retained in the network, rather than to average over uncertainty about whether a directed edge was present, we used the nonzero-mean coefficients and their confidence intervals for the final network. These estimates and confidence intervals are therefore conditional on the corresponding directed edge being present in the selected network structure.

### 3.2 Final Bayesian network

The final Bayesian network (Fig. 5) comprised seven nodes and 14 directed edges. Bootstrap-averaged standardized path coefficients (β) and their 95% confidence intervals are summarized in Fig. 6 and Table S3. Among the 14 edges, eight showed positive effects (blue) and six showed negative effects (red). The 95% bootstrap confidence intervals excluded zero for 11 edges (solid lines), whereas those for the remaining three edges included zero (dashed lines). These edges were retained because they showed high bootstrap support (edge strength = 0.69–0.91), even though their standardized path coefficients were estimated with greater uncertainty. In Fig. 5, edge color indicates the sign of the standardized path coefficient (blue: positive, red: negative), and color intensity is proportional to its absolute value (|β|), with darker edges indicating larger absolute coefficients. Edge width is proportional to edge strength, and arrowhead size is proportional to direction probability. Numbers above the edges indicate β values, and numbers in parentheses below indicate edge strengths. Asterisks indicate statistically significant edges.

The pathway identified by the final Bayesian network began with a positive effect of NPGO3 on JpChumN (β = 0.39; edge strength = 0.98), followed by a negative effect of JpChumN on RusChumN (β = −0.78; edge strength = 1.00). The 95% confidence intervals for both standardized path coefficients excluded zero (Table S3). The JpChumN → RusChumN edge was inferred without directional constraints and had a direction probability of 0.82 (Fig. 4B). Its standardized path coefficient had the largest absolute magnitude among the edges in the final network. The effect of RusChumN on FL was also negative (β = −0.20; edge strength = 0.97), with a 95% confidence interval that excluded zero. JpChumN had a small positive effect on EggSize (β = 0.11; edge strength = 0.83), whereas RusChumN had a small negative effect on EggSize (β = −0.08; edge strength = 0.67). The 95% confidence intervals for both coefficients excluded zero, although their absolute magnitudes were substantially smaller than those of the principal pathways involving maternal traits.

Among the reproductive traits, female fork length (FL) had positive effects for both FL → Fecund (β = 0.62; edge strength = 1.00) and FL → EggSize (β = 0.54; edge strength = 1.00). The 95% confidence intervals for both coefficients excluded zero (Table S3). The Fecund → EggSize edge had a direction probability of 0.64 despite the absence of directional constraints (Fig. 4B), and its effect was negative (β = −0.55; edge strength = 1.00), with a 95% confidence interval that excluded zero. Direct dependence relationships were also identified for River → FL (β = −0.24; edge strength = 1.00) and River → EggSize (β = 0.20; edge strength = 1.00); the 95% confidence intervals for both coefficients excluded zero. The effect for River → Fecund was small and positive (β = 0.08; edge strength = 0.98), with a 95% confidence interval that narrowly excluded zero. EggSize was the terminal node of the final Bayesian network, receiving direct dependence relationships from maternal traits and serving as the endpoint of pathways linking NPGO3, salmon abundance, and river latitude to reproductive traits.

### 3.3 Mediated effects along selected pathways

The directed effects estimated from the second-stage bootstrap were highly correlated with the corresponding nonzero-mean standardized path coefficients from the first-stage bootstrap (*r* = 0.98), indicating close agreement between the two estimates (Fig. S12). The signs were also consistent for all directed effects except River → Fecund (red dot).

The bootstrap distributions of the second-stage estimates for the 14 directed edges and 8 selected mediated pathways are shown in Fig. 7. For each directed edge, the distribution represents the standardized path coefficient (β) estimated across 1000 bootstrap replicates of the fixed final network structure. For each mediated pathway, the distribution represents the product of the standardized path coefficients along the constituent edges. The corresponding means, standard deviations, and 95% bootstrap confidence intervals are provided in Table S4 and Fig. S13.

**Fig. 7.**
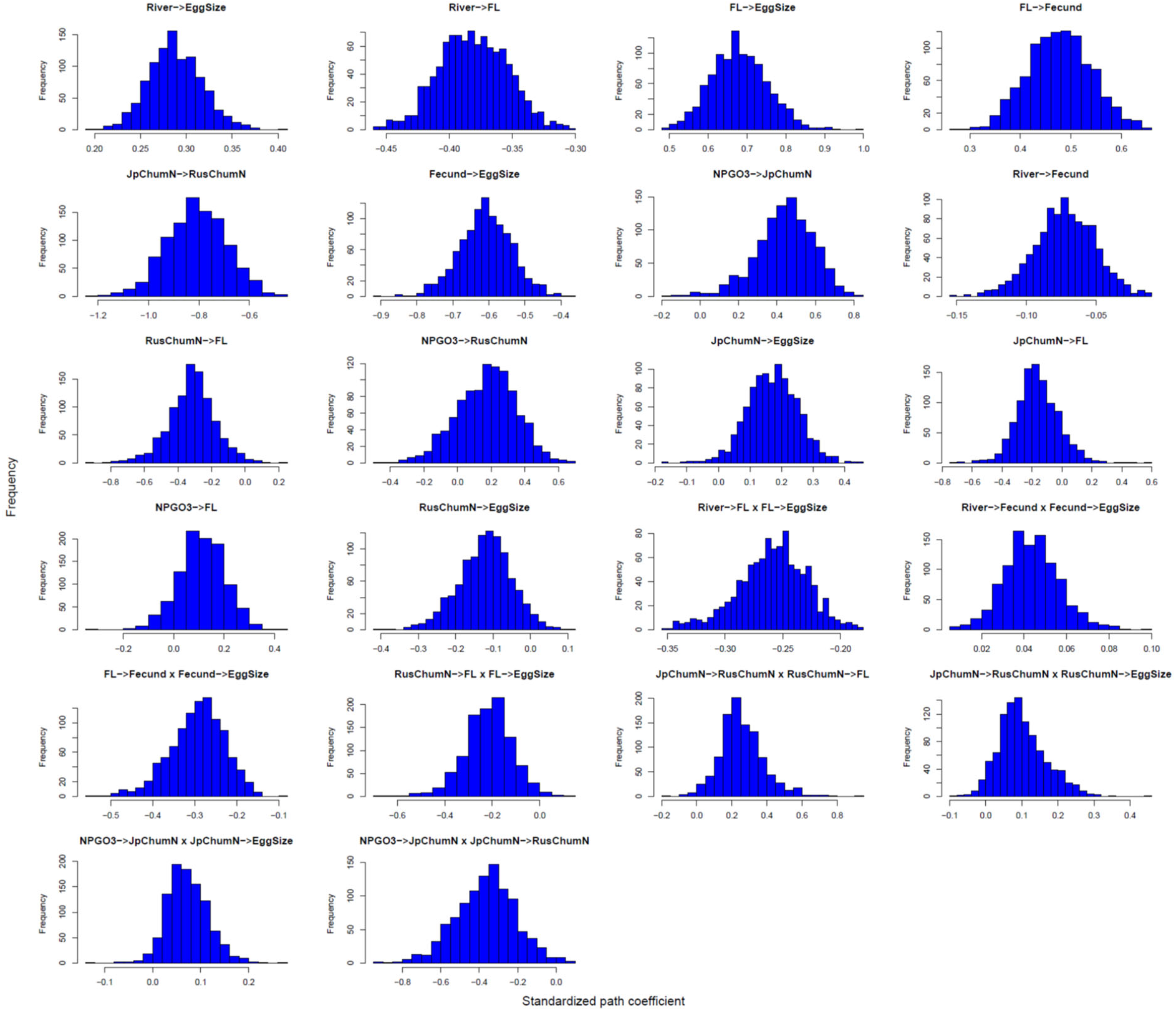
Bootstrap distributions of standardized path coefficients and mediated effects conditional on the final Bayesian network structure (Fig. 5). Histograms show the distributions obtained from 1,000 second-stage bootstrap replicates. For the 14 directed pathways, each histogram represents the bootstrap distribution of the standardized path coefficient for the corresponding retained edge. For the eight mediated pathways, each histogram represents the bootstrap distribution of the product of standardized path coefficients along the constituent edges. The corresponding means and 95% bootstrap confidence intervals are summarized in Table S4.

Six of the eight selected mediated pathways had 95% bootstrap confidence intervals that did not include zero (Table S4, Fig. S13). Negative indirect effects were observed for NPGO3 → JpChumN → RusChumN (β = −0.37), FL → Fecund → EggSize (β = −0.29), River → FL → EggSize (β = −0.26), and RusChumN → FL → EggSize (β = −0.22). Positive indirect effects were observed for JpChumN → RusChumN → FL (β = 0.26) and River → Fecund → EggSize (β = 0.04). In contrast, the mediated effects of JpChumN → RusChumN → EggSize (β = 0.10) and NPGO3 → JpChumN → EggSize (β = 0.07) were small, and their 95% bootstrap confidence intervals included zero.

## 4. Discussion

### 4.1 Oceanographic pathway linking salmon abundance and maternal traits

A previous study documented synchronous long-term declines in egg size, return rate, and Japanese Chum Salmon (JCS) abundance, providing convergent evidence of declining population fitness (Kitada 2026). In the present study, our bootstrap-supported Bayesian network identified a pathway from NPGO3 through JpChumN to RusChumN, followed by a directed effect of RusChumN on female fork length (FL), which was in turn related to egg size. The estimated indirect effect was both negative for NPGO3 → JpChumN → RusChumN (standardized mediated effect, β = −0.37) and RusChumN → FL → EggSize (β = −0.22). Together, these results suggest a pathway in which lower NPGO3 values are linked to lower JCS abundance, followed by higher RCS abundance, smaller female FL, and smaller egg size in JCS.

The negative effect of JpChumN on RusChumN is particularly noteworthy. Among the directed edges in the final network, JpChumN → RusChumN had the largest absolute standardized path coefficient (β = −0.79), and its direction was recovered in 82% of bootstrap replicates without directional constraints. This pattern is consistent with the possibility that declining JCS abundance may have relaxed ecological constraints on RCS, allowing RCS abundance to increase. However, the Bayesian network does not establish that a competitive mechanism underlies this relationship.

Moreover, RCS catches declined after 2015 despite the continued decline in JCS catches (Fig. S14). This period coincided with documented marine heatwave events in the North Pacific (Carvalho et al. 2021), suggesting that unfavorable ocean conditions may have contributed to the decline in RCS catches. However, despite both regions bordering the southern Sea of Okhotsk and thus being expected to experience broadly similar ocean conditions, catches in Eastern Sakhalin increased sharply while those in the Kuriles declined sharply from 2022 onward, despite substantial increases in hatchery releases in both regions (Fig. S7). These contrasting regional trends suggest that unfavorable ocean conditions alone may not adequately explain the observed patterns. Other factors may also have contributed to regional variation in RCS catches, although their relative contributions cannot be determined from the present analysis.

### 4.2 Maternal body size and reproductive traits

Among the reproductive traits, the largest absolute standardized path coefficient was for FL → Fecund (β = 0.62). Female fork length (FL) also had a positive effect on egg size (β = 0.54), whereas fecundity had a negative direct relationship of similar magnitude (β = −0.55). Thus, the positive direct effect of FL on egg size co-occurred with a negative indirect effect of FL on egg size mediated through fecundity (β = −0.29). The position of EggSize at the end of the network highlights how reproductive traits were linked not only to maternal body size and fecundity but also to climate-related oceanographic variability and salmon abundance. These results are consistent with the well-established importance of maternal body size in shaping reproductive traits and offspring characteristics in salmonids (Einum and Fleming 1999). Larger females generally produce larger eggs with greater energy reserves, which can enhance embryonic development, early growth, and survival under variable environmental conditions (Einum and Fleming 1999, 2000).

The Fecund → EggSize direction was recovered in 64% of bootstrap replicates without directional constraints, indicating some uncertainty in the direction of this edge. Nevertheless, the second-stage bootstrap supported a negative mediated effect of FL on egg size through fecundity (Fig. 7, Table S4), indicating that the pathway through fecundity contributed to the effect of maternal body size on egg size. These direct dependence relationships are consistent with the well-established trade-off between offspring size and number in salmonids (Heath et al. 2003) and are biologically plausible given potential constraints on maternal energy allocation (Beacham and Murray 1993; Einum and Fleming 2000; Heath et al. 1999, 2003). Variation in female body size may therefore effects on egg size through both direct and fecundity-mediated pathways. Egg size is also associated with fry size, marine survival, and recruitment success in salmonids (Einum and Fleming 1999, 2000). Together with the findings of Kitada (2026), which documented synchronous long-term declines in JCS abundance, return rate, and egg size, these results suggest that variation in maternal reproductive traits may be relevant to the long-term decline of JCS.

### 4.3 Abundance-mediated variation in egg size

JpChumN had a small positive direct effect on egg size (β = 0.11), whereas RusChumN had a small negative direct effect (β = −0.08). Although both directed edges were statistically supported, their standardized path coefficients were substantially smaller than those for the principal maternal pathways (Fig. 5). Moreover, the mediated effects of JpChumN → RusChumN → EggSize (β = 0.10) and of NPGO3 → JpChumN → EggSize (β = 0.07) were small, and not statistically supported (Table S4). In contrast, the mediated effect of RusChumN → FL → EggSize was statistically supported and negative (β = −0.22). Thus, the results provide stronger support for an effect of abundance on egg size mediated through maternal body size than for a direct effect or a mediated pathway involving abundance alone. The negative indirect effect RusChumN → FL → EggSize is consistent with the possibility that variation in RCS abundance is linked to JCS reproductive traits partly through maternal body size. One possible explanation for the long-term decline in egg size is that climate-related environmental variability and associated changes in salmon abundance may have influenced maternal energy allocation. Importantly, the persistence of the declining trend in egg size after statistically controlling for female body size in the related analysis suggests that changes in egg size cannot be explained by changes in maternal body size alone (Kitada 2026). This finding is consistent with the possibility that changes in reproductive allocation in JCS have contributed to the long-term decline in egg size.

As ectotherms, fishes experience higher metabolic demands at elevated temperatures (Powers et al. 1991; Somero 2010, 2022), potentially reducing the energy available for somatic growth. Such changes in energy allocation could contribute to variation in maternal body size and the amount of energy allocated to individual eggs, thereby influencing egg size. Genetic differentiation among JCS populations may also contribute to geographic variation in energy allocation. Allele frequencies at several SNP loci, including the growth hormone locus Oke_GHII-3129 and the composite mitochondrial DNA haplotypes (mtDNA3) derived from three loci (Oke_Cr30, Oke_Cr386, and Oke_ND3-69; Seeb et al. 2011), show distinctive geographic patterns, with JCS populations deviating markedly from the geographic patterns observed in North Pacific populations (Kitada and Kishino 2021; Fig. S15). However, whether these genetic differences reflect local adaptation, historical population structure, or effects of hatchery practices cannot be determined from the available evidence. Their potential relevance to energy allocation between somatic growth and reproduction also remains unknown.

### 4.4 Geographic variation in maternal traits

Our Bayesian network identified direct pathways from river location (River) to female fork length (FL) and egg size, despite the absence of statistically significant correlations between River and these traits at the river level. The correlations between River and FL (*r* = −0.46), River and fecundity (*r* = −0.23), and River and egg size (*r* = 0.23) were not statistically significant (Fig. S2). Nevertheless, the three northernmost rivers—Tokushibetsu, Ishikari, and Nishibetsu (#1–3 in Fig. S2)—deviated from the main geographic pattern. Among the remaining rivers, egg size showed a strong geographic cline (*r* = 0.93). This geographic variation is consistent with an earlier observation that Chum Salmon from the Tokachi River (#4) produced the largest eggs, whereas those from the Ishikari River (#2) produced the smallest, a pattern reported as early as the 1940s (Watanabe 1955).

Hatchery enhancement of Japanese Chum Salmon (JCS) has continued for more than 140 years, equivalent to approximately 33 generations assuming a generation time of 4.2 years (Kitada 2026). Hatchery fish are released in most large rivers, and escapement of adults for natural spawning is not generally practiced (Kitada 2014). Nevertheless, natural spawning has been observed in rivers in Hokkaido (Miyakoshi et al. 2012), as well as in rivers along the Sea of Japan coast of Honshu (Iida et al. 2018). For semelparous species such as Chum Salmon, Matsuishi et al. (1995) predicted that, when the relative fitness of hatchery fish is 0.6 that of wild fish, more than 99% of wild genes at a biallelic locus would be replaced by hatchery-derived genes within 14 generations, even at a stocking rate of 0.4 (Fig. 5 in Kitada et al. 2019). Gene flow among populations (Kitada and Kishino 2021) may further facilitate the spread of hatchery-derived genes. Thus, most returning JCS may be hatchery fish or wild-born descendants of hatchery fish (Kitada 2020). Consequently, the present study does not directly compare predominantly hatchery-supported and wild populations; rather, it examines variation in reproductive traits among JCS populations that are overwhelmingly hatchery-supported.

Although egg size declined over the study period across the rivers, substantial differences in mean egg size among rivers persisted (Kitada 2026). The persistence of these differences over a long history of hatchery enhancement suggests that geographic variation in egg size has not been eliminated. Egg size is an important reproductive trait with implications for offspring fitness, population productivity, and adaptation to environmental conditions (Quinn et al. 1995; Chambers and Leggett 1996; Mousseau and Fox 1998). However, the present analysis cannot determine whether these differences reflect hatchery selection, environmental variation, genetic differentiation, or interactions among these factors.

### 4.5 Ecological assumptions of our Bayesian network modeling

We represented marine environmental conditions using the NPGO index in the release year (NPGO3), which was assumed to represent conditions experienced during the first year of marine life (ages 0–1). However, Japanese Chum Salmon (JCS) abundance (JpChumN) is influenced by environmental conditions encountered throughout the marine phase, spanning approximately 2–6 years (Fig. 1). Incorporating environmental variables with multiple time lags into the Bayesian network would substantially increase model complexity and introduce strong correlations among NPGO variables, potentially reducing the stability and interpretability of network inference. It would also require more explicit matching of environmental conditions to different age classes and marine life-history stages. We therefore used NPGO3 as a parsimonious proxy for broad-scale marine environmental variability. Although this approach simplifies the system, it may be biologically reasonable because mortality can be particularly high during the first year at sea (Beamish and Mahnken 2001), and substantial mortality of hatchery-released fry has been suggested in Japanese coastal waters (Kitada et al. 2025). These observations suggest that environmental conditions during this period may have disproportionate effects on subsequent survival and abundance.

The distributions of Chum, Pink (*O. gorbuscha*), and Sockeye Salmon (*O. nerka*) overlap extensively across the North Pacific (Langan et al. 2024), and these species exhibit similar diet compositions, with diet overlap estimated at 32% between Chum and Pink Salmon and 31% between Chum and Sockeye Salmon (Graham et al. 2021). Because Pink Salmon complete their marine life within approximately 18 months (NPAFC 2026b), multiple Russian Pink Salmon year classes may overlap temporally with a single JCS cohort and potentially influence its return rate (Kitada et al. 2025). We therefore examined Russian Pink Salmon catch in number (RusPinkN) as an additional potential correlate of JCS abundance in the Bayesian network. JpChumN and RusPinkN were negatively correlated (*r* = −0.43), whereas RusChumN and RusPinkN were positively correlated (*r* = 0.44) (Fig. S16). We also examined Russian Sockeye Salmon abundance, but its strong correlation with RusChumN (*r* = 0.90) indicated substantial redundancy. Moreover, Sockeye Salmon have a more complex life history, typically using lakes as rearing habitats for one to three years before entering the ocean and spending one to four years in the ocean before returning to freshwater to spawn (NPAFC 2026b). We therefore did not include Sockeye Salmon abundance as a separate node in the network.

When RusPinkN was added to the Bayesian network (Fig. 8), 19 of the 32 directed edges examined met both criteria of edge strength >0.6 and direction probability >0.5 (Fig. S17). Of these 19 edges, nine had positive and 10 had negative standardized path coefficients. Using the nonzero-mean approach, the 95% confidence intervals for the mean standardized path coefficients excluded zero for 13 edges (solid lines), whereas those for the remaining six edges included zero (dashed lines). The relationships among the other seven nodes (Table 1) remained unchanged from the network without RusPinkN (Fig. 5). RusPinkN received directed edges from NPGO3 and JpChumN with negative effects and from RusChumN with a positive effect. None of these effects was statistically supported, although the estimated standard path coefficients indicated that RusPinkN tended to increase with RusChumN and decrease with JpChumN (Fig. 8). These patterns are compatible with ecological interactions among salmon populations whose abundances changed concurrently across the North Pacific (Fig. S16), but the present analysis cannot identify the underlying mechanisms.

**Fig. 8.**
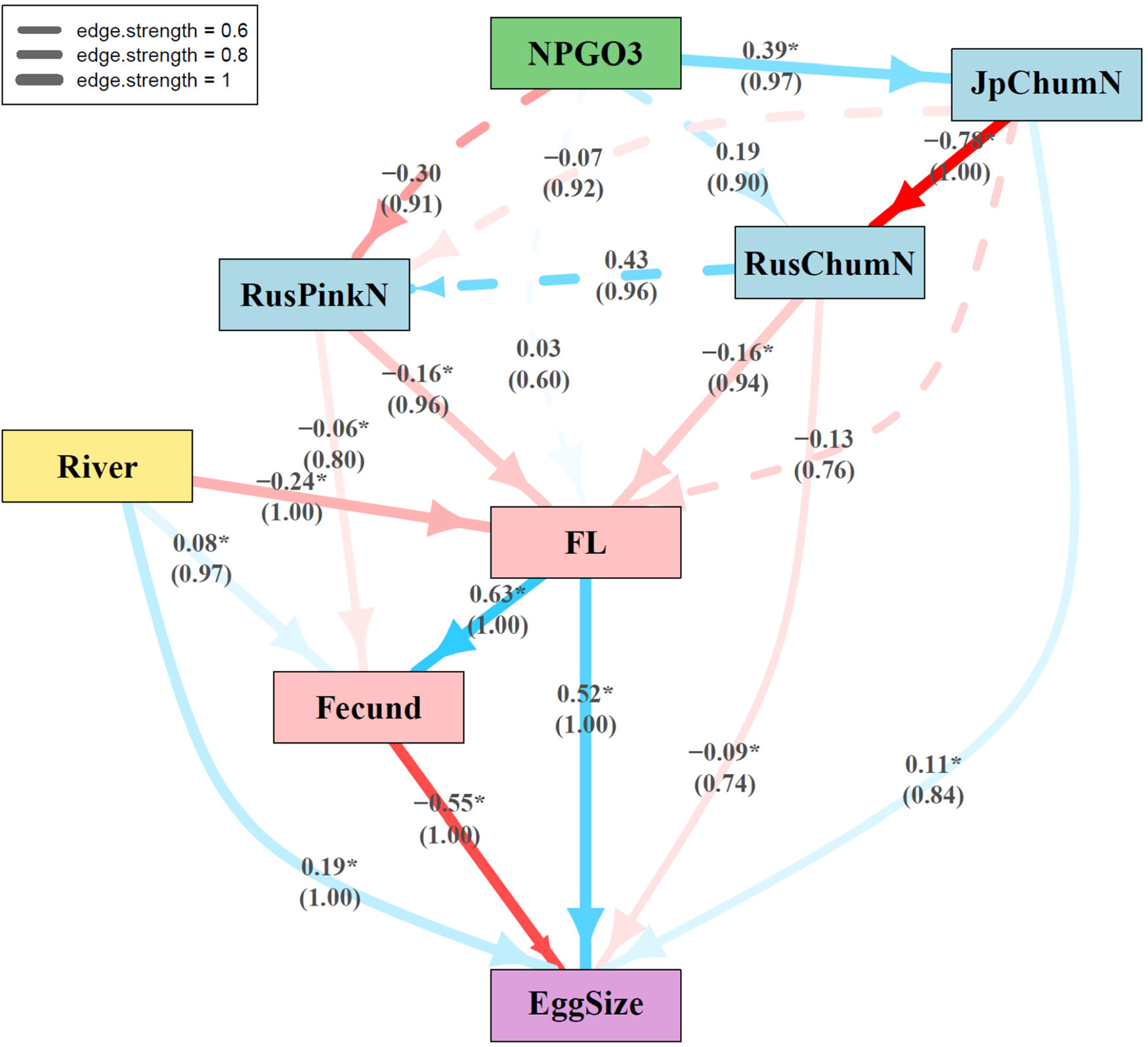
Final bootstrap-supported Bayesian network incorporating Russian pink salmon abundance, inferred from 1,000 block bootstrap replicates in which years (1999–2019) were resampled while retaining all individuals within each year (*n* = 12,428), thereby accounting for temporal dependence. Edges with edge strength > 0.6 and direction probability > 0.5 were retained for construction of the final Bayesian network, resulting in a network with eight nodes and 19 directed edges (see Fig. S16). The width of each edge is proportional to edge strength, and the size of the arrowhead is proportional to direction probability. Numbers shown above the edges are standardized path coefficients, and numbers in parentheses below indicate edge strengths. Asterisks indicate edges whose bootstrap 95% confidence intervals for standardized path coefficients excluded zero (see Fig. S17). Dashed lines indicate edges whose bootstrap 95% confidence intervals for standardized path coefficients included zero. Color intensity is proportional to absolute standardized path coefficient.

RusPinkN also had a negative effect on female fork length (FL), with an estimated standardized path coefficient of β = −0.16 (edge strength = 0.96), similar in magnitude to that for RusChumN (β = −0.16; edge strength = 0.94) (Fig. 8, Table S5, Fig. S18). This direct dependence relationship indicates that Russian Pink Salmon abundance had a negative effect on JCS female body size. However, because the incoming edges to RusPinkN were not statistically supported, the expanded network does not provide strong evidence for a pathway linking Russian Pink Salmon abundance to the decline in female body size in JCS. This result is therefore best interpreted as a supplementary association rather than as evidence of a specific ecological mechanism.

These results should also be interpreted in light of the temporal ordering of the abundance indices. The decline in JCS abundance preceded the subsequent increases in RCS and Russian Pink Salmon abundance. Thus, the network results do not indicate that increases in Russian Chum or Pink Salmon abundance preceded or explained the initial decline in JCS. Rather, the observed temporal patterns are consistent with the possibility that changes in Russian salmon populations occurred subsequently in association with broader changes in the North Pacific ecosystem.

### 4.6 Statistical assumptions in our Bayesian network modeling

To assess the functional form of relationships among the variables, we examined pairwise scatter plots and correlation plots for the six variables in the final Bayesian network other than River, together with Russian Pink Salmon abundance (RusPinkN) and mean age at return. The plots showed generally weak or moderate linear associations, with no pronounced curvilinear patterns (Fig. S19). Mean age at return showed relatively small and non-significant correlations with the other seven variables (Fig. S20). Mean age was included in this supplementary assessment because age at maturity is an important life-history characteristic and showed substantial temporal variation (Fig. 5 in Kitada 2026). In that study, the decline in mean age at return became apparent only after approximately 2010, whereas a marked change in the rate of abundance decline occurred around 2005, indicating that the temporal patterns of age composition and abundance decline were not closely aligned. Although mean age was therefore not included as a node in the Bayesian network, its weak correlations with the other variables provided an additional check on potential confounding by age composition. These checks provided no indication that pronounced non-linear relationships were a major concern for the fitted Gaussian Bayesian network.

Because temporal dependence and synchronous variation among rivers reduce the effective sample size relative to the number of observed individuals, we performed block bootstrap resampling in which years were resampled while retaining all individuals within each year. As a sensitivity analysis, we also examined a Bayesian network inferred without accounting for temporal dependence (Fig. S21). This network had the same structure as the final bootstrap-supported network (Fig. 5), with the same seven nodes and 14 directed edges. However, all retained edges had extremely high edge strengths (≥0.97; Fig. S10A). This uniformly high support may have resulted from treating observations as independent despite temporal dependence and synchronous variation among rivers, thereby overstating the amount of independent information available and underestimating uncertainty. Although the network recovered the same 14 directed edges, the direction of the edge between JpChumN and RusChumN was less certain, with the direction probability approaching 0.5 (Fig. S10). Point estimates of the standardized path coefficients were very similar between the two approaches (*r* = 0.995), whereas the analysis that did not account for temporal dependence yielded substantially smaller standard errors and narrower confidence intervals (Figs. S22, S23). These results suggest that ignoring temporal dependence and river-level correlation may lead to underestimation of uncertainty, even when the estimated network structure and path coefficients remain largely unchanged.

Taken together, these findings complement the long-term declines in egg size, return rate, and abundance reported for Japanese Chum Salmon (JCS) by Kitada (2026). The persistence of the egg-size decline after statistically controlling for maternal body size suggests that body size alone does not fully explain this long-term trend. Our Bayesian network analysis identified a directed pathway linking lower NPGO3 values to lower JCS abundance, followed by higher Russian Chum Salmon (RCS) abundance, smaller female fork length (FL), and smaller egg size in JCS. A separate maternal pathway linked FL to egg size through fecundity. River location also showed direct dependence relationships with FL, fecundity, and egg size, highlighting geographic variation in maternal reproductive traits. The Bayesian network provides an integrated framework for identifying direct dependence relationships among oceanographic variability, salmon abundance indices, and maternal reproductive traits. Further investigation is needed to clarify the mechanisms underlying long-term changes in reproductive traits.

## Author contributions

Conceptualization: SK, HK; Data curation: SK; Formal analysis: SK, HK; Funding acquisition: SK, HK; Methodology: HK; Software: HK, SK; Validation: HK, SK; Visualization: SK, HK; Writing – original draft: SK; Writing – review and editing: HK, SK. All authors approved the final version of the manuscript.

## Acknowledgements

We thank the anonymous Associate Editor and reviewers for their insightful and constructive comments. We also gratefully acknowledge Marco Scutari for his valuable guidance on Bayesian network analysis and the use of the `bnlearn` package. We thank the many biologists, fishery and hatchery managers, and fishers who contributed to the long-term monitoring and data collection efforts underlying this study. We are also grateful to the Fisheries Research and Education Agency (FRA) and the North Pacific Anadromous Fish Commission (NPAFC) for compiling and maintaining the publicly available databases that made this analysis possible. ChatGPT (OpenAI, GPT-5.5) was used to assist with English editing, manuscript revision, and code troubleshooting. The authors take full responsibility for all scientific interpretations, analyses, and conclusions. This work was supported by the Japan Society for the Promotion of Science (JSPS) through Grants-in-Aid for Scientific Research (KAKENHI) (Nos. 18K05781 to SK and 22K11950 to HK).

## Data availability statement

The data generated or analyzed in this study are fully described in the published article and its Supplementary Information. Analysis scripts and data will be available on acceptance.

## References

Amoroso, R. O., Tillotson, M. D., and Hilborn, R. 2017. Measuring the net biological impact of fisheries enhancement: Pink salmon hatcheries can increase yield, but with apparent costs to wild populations. Can. J. Fish. Aquat. Sci. 74(8): 1233–1242. doi10.1139/cjfas-2016-0334

Araki, H., Cooper, B., and Blouin, M. S. 2007. Genetic effects of captive breeding cause a rapid, cumulative fitness decline in the wild. Science 318: 100–103. doi10.1126/science.11456

Beacham, T. D., and Murray, C. B. 1993. Fecundity and egg size variation in North American Pacific salmon (*Oncorhynchus*). J. Fish Biol. 42(4): 485–508. doi10.1111/j.1095-8649.1993.tb00354.x

Beamish, R. J., and Mahnken, C. 2001. A critical size and period hypothesis to explain natural regulation of salmon abundance and the linkage to climate and climate change. Prog. Oceanogr. 49(1–4): 423–437. doi10.1016/S0079-6611(01)00034-9

Carvalho, K. S., Smith, T. E., and Wang, S. E. 2021. Bering Sea marine heatwaves: Patterns, trends and connections with the Arctic. J. Hydro. 600: 126462. doi10.1016/j.jhydrol.2021.126462.

Chambers, R. C., and Leggett, W. C. 1996. Maternal influences on variation in egg sizes in temperate marine fishes. Am. Zoolog. 36(2): 180–196. doi10.1093/icb/36.2.180

Cheung, W. W., Frölicher, T. L., Lam, V. W., Oyinlola, M. A., Reygondeau, G., Sumaila, U. R., … and Wabnitz, C. C. 2021. Marine high temperature extremes amplify the impacts of climate change on fish and fisheries. Science Advances, 7(40): eabh0895. doi10.1126/sciadv.abh089

Cheung, W. W., Pauly, D., and Sumaila, U. R. 2025. Hope or despair revisited: assessing progress and new challenges in global fisheries. Fish Fish. 26(2): 257–269. doi10.1111/faf.12877

Cline, T. J., Ohlberger, J., and Schindler, D. E. 2019. Effects of warming climate and competition in the ocean for life-histories of Pacific salmon. Nature Ecol. Evol. 3(6): 935–942. doi10.1038/s41559-019-0901-7

Christie, M. R., Marine, M. L., French, R. A., and Blouin, M. S. 2012. Genetic adaptation to captivity can occur in a single generation. Proc. Natl. Acad. Sci.109(1): 238–242. doi10.1073/pnas.111107310

Christie, M. R., Ford, M. J., and Blouin, M. S. 2014. On the reproductive success of early-generation hatchery fish in the wild. Evol. Appl. 7(8): 883–896. doi10.1111/eva.12183

Crozier, L. G., Hendry, A. P., Lawson, P. W., Quinn, T. P., Mantua, N. J., Battin, J., … and Huey, R. 2008. Potential responses to climate change in organisms with complex life histories: evolution and plasticity in Pacific salmon. Evol. Appl. 1(2): 252–270. doi10.1111/j.1752-4571.2008.00033.x

Crozier, L. G., Burke, B. J., Chasco, B. E., Widener, D. L., and Zabel, R. W. 2021. Climate change threatens Chinook salmon throughout their life cycle. *Comm*. Biol. 4(1): 222. doi10.1038/s42003-021-01734-w

Csardi, G., and Nepusz, T. (2006). The igraph software. Complex syst, 1695: 1–9.

Cunningham, C. J., Westley, P. A., and Adkison, M. D. 2018. Signals of large scale climate drivers, hatchery enhancement, and marine factors in Yukon River Chinook salmon survival revealed with a Bayesian life history model. Global Change Biol. 24(9), 4399– 4416. doi10.1111/gcb.14315

Davis, M. J., Anthony, J., Ward, E. J., Firman, J., and Lorion, C. 2023. Coherence among Oregon Coast coho salmon populations highlights increasing relative importance of marine conditions for productivity. Fish. Ocean. 32(3): 293–310. doi10.1111/fog.12630

Debertin, A. J., Irvine, J. R., Holt, C. A., Oka, G., and Trudel, M. 2017. Marine growth patterns of southern British Columbia Chum Salmon explained by interactions between density-dependent competition and changing climate. Can. J. Fish. Aquat. Sci. 74(7): 1077– 1087. doi10.1139/cjfas-2016-0265

Di Lorenzo, E., Schneider, N., Cobb, K. M., Franks, P. J. S., Chhak, K., Miller, A. J., … and Rivière, P. 2008. North Pacific Gyre Oscillation links ocean climate and ecosystem change. Geophysical Res. Let. 35(8): L08607. doi:10.1029/2007GL032838. doi.org/10.1029/2007GL032838

Dunmall, K. M., McNicholl, D. G., Zimmerman, C. E., Gilk-Baumer, S. E., Burril, S., and von Biela, V. R. 2022. First juvenile Chum Salmon confirms successful reproduction for Pacific salmon in the North American Arctic. Can. J. Fish. Aquat. Sci. 79(5): 703–707. doi10.1139/cjfas-2022-0006

Einum, S., and Fleming, I. A. 1999. Maternal effects of egg size in brown trout (*Salmo trutta*): Norms of reaction to environmental quality. Proc. Royal Soc. Lond. B 266: 2095–2100. doi10.1098/rspb.1999.0893

Einum, S., and Fleming, I. A. 2000. Highly fecund mothers sacrifice offspring survival to maximize fitness. Nature 405: 565–567. doi10.1038/35014600

Felsenstein, J. 1985. Confidence limits on phylogenies: an approach using the bootstrap. Evolution, 39(4): 783–791. doi10.1111/j.1558-5646.1985.tb00420.x

Fleming, I. A., and Gross, M. R. 1990. Latitudinal clines: A trade-off between egg number and size in Pacific salmon. Ecology 71: 1–11. doi10.2307/1940241

Ford, M. J., Lindley, S. T., Barnas, K. A., Shelton, A. O., Spence, B. C., Weitkamp, L. A., … and Williams, T. H. 2025. Abundance trends of Pacific salmon during a quarter century of ESA protection. Fish Fish. 26(6): 1087–1106. doi10.1111/faf.70019

FRA (Japan Fisheries Research and Education Agency). 2026. Salmon Data Base. FRA. (in Japanese). www.fra.go.jp/shigen/salmon/sdb.html. [Accessed February, 2026]

Graham, C., Pakhomov, E. A., and Hunt, B. P. 2021. Meta-analysis of salmon trophic ecology reveals spatial and interspecies dynamics across the North Pacific Ocean. Front. Mar. Sci. 8, 618884. doi10.3389/fmars.2021.618884

Heath, D. D., Fox, C. W., and Heath, J. W. 1999. Maternal effects on offspring size: variation through early development of chinook salmon. Evolution 53(5): 1605–1611. doi10.1111/j.1558-5646.1999.tb05424.x

Heath, D. D., Heath, J. W., Bryden, C. A., Johnson, R. M., and Fox, C. W. 2003. Rapid evolution of egg size in captive salmon. Science 299(5613): 1738–1740. doi10.1126/science.107970

Hilborn, R., and Eggers, D. 2000. A review of the hatchery programs for pink salmon in Prince William Sound and Kodiak Island, Alaska. Trans. Am. Fish. Soc. 129(2): 333–350. doi.org/10.1577/1548-8659(2000)129<0333:AROTHP>2.0.CO;2

Hilborn, R., Quinn, T. P., Schindler, D. E., and Rogers, D. E. 2003. Biocomplexity and fisheries sustainability. Proc. Natl. Acad. Sci. 100(11), 6564–6568.doi10.1073/pnas.103727410

Iida, M., Yoshino, K., and Katayama, S. 2018. Current status of natural spawning of Chum Salmon *Oncorhynchus keta* in rivers with or without hatchery stocking on the Japan Sea side of northern Honshu, Japan. Fish. Sci. 84(3): 453–459. doi10.1007/s12562-018-1192-7

Jeffrey, K. M., Côté, I. M., Irvine, J. R., and Reynolds, J. D. 2017. Changes in body size of Canadian Pacific salmon over six decades. Can. J. Fish. Aquat. Sci. 74(2): 191–201. doi10.1139/cjfas-2015-060

Kitada, S. 2014. Japanese chum salmon stock enhancement: current perspective and future challenges. Fish. Sci. 80(2): 237–249. doi10.1007/s12562-013-0692-8

Kitada, S. 2018. Economic, ecological and genetic impacts of marine stock enhancement and sea ranching: A systematic review. Fish Fish. 19: 511–532. doi10.1111/faf.12271

Kitada, S. 2020. Lessons from Japan marine stock enhancement and sea ranching programmes over 100 years. Rev. Aquacult. 12: 1944–1969. doi10.1111/raq.12418

Kitada, S. 2026. Long-term decline in egg size of Japanese hatchery Chum Salmon links to declines in return rate and abundance. Can. J. Fish. Aquat. Sci. 83. doi10.1139/cjfas-2026-0158.

Kitada, S., and Kishino, H. 2021. Population structure of Chum Salmon and selection on the markers collected for stock identification. Ecol. Evol. 11(20): 13972–13985. doi10.1002/ece3.8102

Kitada, S., Myers, K. W., and Kishino, H. 2025. Hatcheries to high seas: climate change connections to salmon marine survival. Ecol. Evol. 15(6): e71504. doi10.1002/ece3.71504

Kitada, S., Nakajima, K., Hamasaki, K., Shishidou, H., Waples, R. S., and Kishino, H. 2019. Rigorous monitoring of a large-scale marine stock enhancement program demonstrates the need for comprehensive management of fisheries and nursery habitat. Sci. Rep. 9(1): 5290. doi10.1038/s41598-019-39050-3

Langan, J. A., Cunningham, C. J., Watson, J. T., and McKinnell, S. 2024. Opening the black box: new insights into the role of temperature in the marine distributions of Pacific salmon. Fish Fish. 25(4): 551–568. doi10.1111/faf.12825

Lewis, B., Grant, W. S., Brenner, R. E., and Hamazaki, T. 2015. Changes in size and age of Chinook salmon *Oncorhynchus tshawytscha* returning to Alaska. PLoS One 10(6): e0130184. doi10.1371/journal.pone.0130184

Malick, M. J., Losee, J. P., Marston, G., Agha, M., Berejikian, B. A., Beckman, B. R., and Cooper, M. 2023. Fecundity trends of Chinook salmon in the Pacific Northwest. Fish Fish. 24: 454–465. doi10.1111/faf.12738

MAFF (Ministry of Agriculture, Forestry and Fisheries). 2026a. Annual Statistics of Fisheries and Aquaculture Production. MAFF. (in Japanese). Accessed September 2, 2026. https://www.maff.go.jp/j/tokei/kouhyou/kensaku/bunya6.html

MAFF (Ministry of Agriculture, Forestry and Fisheries). 2026b. Fishery Census. MAFF. (in Japanese). Accessed October 11, 2026. https://www.maff.go.jp/j/tokei/census/fc/index.html

Matsuishi, T., Kishino, H., and Numachi, K. 1995. A model of gene displacement by stocking activities. Nippon Suisan Gakkaishi 61: 326–330. (in Japanese with English abstract) doi.org/10.2331/suisan.61.326

Max Planck Institute. 2023. Climate Data Operators. Hamburg: Max Planck Institute für Meteorologie. code.mpimet.mpg.de/projects/cdo (accessed 14 June 2023).

Miyakoshi, Y., Urabe, H., Saneyoshi, H., Aoyama, T., Sakamoto, H., Ando, D., and Nagata, M. 2012. The occurrence and run timing of naturally spawning Chum Salmon in northern Japan. Environ. Biol. Fishes 94(1): 197–206. doi10.1007/s10641-011-9872-5

Miyakoshi, Y., Nagata, M., Kitada, S., and Kaeriyama, M. 2013. Historical and current hatchery programs and management of Chum Salmon in Hokkaido, northern Japan. Rev. Fish. Sci. 21(3-4): 469–479. doi10.1080/10641262.2013.836446

Morita, K., Saito, T., Miyakoshi, Y., Fukuwaka, M. A., Nagasawa, T., and Kaeriyama, M. 2006. A review of Pacific salmon hatchery programmes on Hokkaido Island, Japan. ICES J. Mar. Sci. 63(7): 1353–1363. doi10.1016/j.icesjms.2006.03.024

Mousseau, T. A., and Fox, C. W. 1998. The adaptive significance of maternal effects. Trends Ecol. Evol 13(10): 403–407. doi10.1016/S0169-5347(98)01472-4

Naish, K. A., Taylor III, J. E., Levin, P. S., Quinn, T. P., Winton, J. R., Huppert, D., and Hilborn, R. 2007. An evaluation of the effects of conservation and fishery enhancement hatcheries on wild populations of salmon. Adv. Mar. Biol. 53: 61–194. doi10.1016/S0065-2881(07)53002-6

NOAA (National Oceanic and Atmospheric Administration). 2026. NOAA Physical Sciences Laboratory, psl.noaa.gov/data/timeseries/month/DS/NPGO/ [accessed May 2026].

NPAFC (North Pacific Anadromous Fish Commission) 2026a. NPAFC statistics: Pacific salmonid catch and hatchery release data. Vancouver: North Pacific Anadromous Fish Commission. www.npafc.org/statistics/ [accessed August 2025].

NPAFC (North Pacific Anadromous Fish Commission) 2026b. *Species*. Vancouver: North Pacific Anadromous Fish Commission. www.npafc.org/species/ [accessed June 2026].

Ohlberger, J., Cline, T. J., Schindler, D. E., and Lewis, B. 2023. Declines in body size of sockeye salmon associated with increased competition in the ocean. Proc. Royal Soc. Lond. B 290: 20222248. doi10.1098/rspb.2022.2248

Oke, K. B., Cunningham, C. J., Westley, P. A. H., Baskett, M. L., Carlson, S. M., Clark, J., … and Palkovacs, E. P. 2020. Recent declines in salmon body size impact ecosystems and fisheries. Nature Comm. 11: 4155. doi10.1038/s41467-020-17726-z

Pearl, J. 1985. Bayesian networks: A model of self-activated memory for evidential reasoning. In Proceedings of the annual meeting of the Cognitive Science Society (Vol. 7). https://escholarship.org/uc/item/0vr7830n

Powers, D. A., Lauerman, T., Crawford, D., and DiMichele, L. 1991. Genetic mechanisms for adapting to a changing environment. Ann. Rev. Genet. 25: 629–660. 10.1146/annurev.ge.25.120191.003213

Quinn, T. P., Hendry, A. P., and Wetzel, L. A. 1995. The influence of life history trade-offs and the size of incubation gravels on egg size variation in sockeye salmon (*Oncorhynchus nerka*). Oikos: 425–438. doi10.2307/3545987

R Core Team 2026. R: A language and environment for statistical computing. R Foundation for Statistical Computing.

Riddell, B. E., Pearsall, I., and Rosenberger, A. 2024. A review of Pacific salmon hatcheries in British Columbia, Canada, and interactions with natural populations. Fisheries 49(7): 303–318. doi10.1002/fsh.11091

Ruggerone, G. T., and Irvine, J. R. 2018. Numbers and biomass of natural-and hatchery-origin pink salmon, Chum Salmon, and sockeye salmon in the North Pacific Ocean, 1925–2015. Mar. Coast. Fish. 10(2): 152–168. doi10.1002/mcf2.10023

Scutari, M. 2010. Learning Bayesian networks with the bnlearn R package. Journal of Statistical Software 35: 1–22. doi10.18637/jss.v035.i03

Scutari, M., and Denis, J. B. 2021. Bayesian networks: with examples in R. CRC press. doi.org/10.1201/9780429347436

Seeb, L. W., Templin, W. D., Sato, S., Abe, S., Warheit, K., Park, J. Y., and Seeb, J. E. 2011. Single nucleotide polymorphisms across a species’ range: implications for conservation studies of Pacific salmon. Molec. Ecol. Resour. 11: 195–217. doi10.1111/j.1755-0998.2010.02966.x

Somero, G. N. 2010. The physiology of climate change: how potentials for acclimatization and genetic adaptation will determine ‘winners’ and ‘losers’. J. Exp. Biol. 213: 912–920. 10.1242/jeb.037473

Somero, G. N. 2022. The Goldilocks Principle: A unifying perspective on biochemical adaptation to abiotic stressors in the sea. Ann. Rev. Mar. Sci. 14: 1–23. 10.1146/annurev-marine-022521-102228

Urawa, S., Beacham, T., Sutherland, B., and Sato, S. 2022. Winter distribution of Chum Salmon in the Gulf of Alaska: A review. North Pac. Anad. Fish Comm. Tech. Rep. 18: 83–87. doi10.23849/npafctr18

Waples, R. S. 1991. Genetic interactions between hatchery and wild salmonids: lessons from the Pacific Northwest. Can. J. Fish. Aquat. Sci. 48(S1): 124–133. doi10.1139/f91-311

Waples, R. S., and Drake, J. 2004. Risk/benefit considerations for marine stock enhancement: a Pacific salmon perspective. In Stock Enhancement and Sea Ranching: Developments, Pitfalls and Opportunities, edited by K. M. Leber, S. Kitada, H. L. Blankenship, T. Svåsand, 260–306. Blackwell Publishing.

Waples, R. S., Beechie, T., and Pess, G. R. 2009. Evolutionary history, habitat disturbance regimes, and anthropogenic changes: what do these mean for resilience of Pacific salmon populations? Ecol. Soc. 14(1) 3. jstor.org/stable/26268053

Watanabe, M. 1955. Some observation on the eggs of mature salmon (*Oncorhynchus keta*) in Hokkaido, with special reference to the race of salmon as characterized by the size of their eggs. Sci. Rep. Hokkaido Salmon Hatchery 10(1,2): 7–20. (in Japanese)

