## Supplementary Information for "Climate-driven fitness decline in Japanese Chum Salmon reshapes North Pacific salmon biogeography, with reproductive consequences: A Bayesian network analysis"

This PDF file includes:

Glossary of Bayesian network terms
Tables S1–S5
Figures S1–S23

**Glossary of Bayesian network terms**

The definitions below follow standard terminology used in Bayesian network analysis and the `bnlearn` package (Scutari 2010; Scutari and Denis 2021).

**Arc (Edge).** An edge, or arc, represents a statistical dependency between two nodes in a Bayesian network. In a directed network, the arrow indicates the direction of the dependency.

**Blacklist.** A blacklist is a set of constraints that prevents specified directed edges from being included in the learned network. In this study, blacklist constraints were used to restrict possible edge directions based on a priori biological and ecological knowledge (Domain knowledge) about the relationships among variables.

**Directed acyclic graph (DAG).** A directed acyclic graph is a graph consisting of nodes connected by directed edges, in which no path forms a closed directed cycle. Bayesian network structures are represented as DAGs.

**Direction probability.** Direction probability is the probability that an edge is oriented in a particular direction among bootstrap replicates in which an edge between the two nodes is recovered. In this study, it was calculated as boot.p / (boot.p + opp.boot.p), where boot.p and opp.boot.p are the bootstrap support probabilities for the inferred and opposite directions, respectively.

**Domain knowledge.** Biological, ecological, and other subject-specific knowledge that informs the selection of variables, the specification of constraints on network structure, and the interpretation of inferred relationships. Domain knowledge is a key information in Bayesian network analysis because it helps ensure that the inferred network is biologically plausible and that its structure and relationships are interpreted in the context of established evidence. In this study, domain knowledge comprised the biological and ecological information and supporting evidence described in Sections 2.2–2.5, which informed variable selection, the specification of structural constraints, and the interpretation of the inferred network.

**Edge strength.** Edge strength represents the bootstrap support for the presence of an edge between a pair of nodes, regardless of its direction. In this study, it was calculated as boot.p + opp.boot.p, where boot.p and opp.boot.p are the bootstrap support probabilities for the inferred and opposite directions, respectively.

**Hill climbing.** Hill climbing is a heuristic algorithm used to learn the Bayesian network structure by iteratively making local changes to a network that improve the Bayesian information criterion (BIC) score, which balances model fit and model complexity. The BIC score is used to compare alternative network structures for a given set of nodes and data. The procedure continues until no further change improves the score, subject to the specified structural constraints. BIC values from networks containing different sets of nodes should not be interpreted as indicating which set of variables is more appropriate, because such networks represent probability distributions over different variable spaces and have different parameterizations.

**Node.** A node represents a variable in a Bayesian network. In this study, nodes represent environmental, salmon abundance, life-history, or fitness-related variables.

**Parent and child nodes.** In a directed Bayesian network, a parent node is connected to a child node by an edge directed from the parent to the child. The child node is modelled conditional on its parent node(s).

**References**

Scutari, M. 2010. Learning Bayesian networks with the bnlearn R package. *Journal of Statistical Software* 35: 1–22. doi.org/10.18637/jss.v035.i03

Scutari, M., and Denis, J. B. 2021. *Bayesian networks: with examples in R*. CRC press. doi.org/10.1201/9780429347436

**Table S1.** Geographic coordinates and areas of the 16 chum salmon distribution areas used for SST analysis in this study.

| Num. | Area | Latitude(°N) | Longitude | Area ($km^2$) |
|---|---|---|---|---|
| 1 | Chukchi Sea | 68–71 | 180°E–160°W | 259,775 |
| 2 | Northern Bering Sea | 60–65 | 180°E–165°W | 428,054 |
| 3 | Western Bering Sea | 50–60 | 163°E–180°E | 1,204,090 |
| 4 | Western North Pacific | 42–50 | 158°E–180°E | 1,510,432 |
| 5 | Western Kamchatka | 59–61.8 | 156°E–161°E | 85,493 |
| 6 | Northern Sea of Okhotsk | 51–59 | 145°E–156°E | 623,578 |
| 7 | Russian continental coast | 54–57 | 138°E–145°E | 147,051 |
| 8 | Southern Sea of Okhotsk | 43–51 | 145°E–158°E | 876,262 |
| 9 | Northern Pacific coast of Japan | 36–43 | 142°E–150°E | 533,942 |
| 10 | Northern Sea of Japan | 38–45 | 136°E–140°E | 259,128 |
| 11 | Eastern Bering Sea | 50–60 | 180°E–160°W | 1,416,576 |
| 12 | Eastern North Pacific | 42–50 | 180°E–155°W | 1,716,399 |
| 13 | Gulf of Alaska | 47–59 | 155°W–139°W | 1,082,762 |
| 14 | British Columbia | 49–57 | 139°W–130°W | 535,319 |
| 15 | Washington and Oregon | 40–49 | 135°W–125°W | 792,881 |
| 16 | California | 33–40 | 132°W–123°W | 625,776 |

**Table S2.** Biological constraints imposed on the Bayesian networks, as defined by the blacklist. It was assumed that environmental variables are not influenced by individual traits or salmon abundance, and that salmon abundance is not influenced by individual traits. It was also assumed that egg size and fecundity do not influence female fork length (FL), because both variables represent reproductive traits of the same female and were therefore considered to be determined by maternal body size and condition.

| Num. | From | To |
|---:|---|---|
| 1 | FL | NPGO3 |
| 2 | Fecund | NPGO3 |
| 3 | EggSize | NPGO3 |
| 4 | FL | River |
| 5 | Fecund | River |
| 6 | EggSize | River |
| 7 | FL | JpChumN |
| 8 | Fecund | JpChumN |
| 9 | EggSize | JpChumN |
| 10 | FL | RusChumN |
| 11 | Fecund | RusChumN |
| 12 | EggSize | RusChumN |
| 13 | JpChumN | NPGO3 |
| 14 | RusChumN | NPGO3 |
| 15 | JpChumN | River |
| 16 | RusChumN | River |
| 17 | EggSize | FL |
| 18 | Fecund | FL |

**Table S3.** Bootstrap estimates and 95% confidence intervals for standardized path coefficients of the 14 directed edges retained in the final Bayesian network based on edge strength > 0.6 and direction probability > 0.5. Mean and SD summarize the bootstrap distribution of non-zero standardized coefficients from networks in which the corresponding directed edge was present. The 95% bootstrap confidence intervals were estimated using the normal approximation as mean ± 1.96 SD. Bold values indicate statistically significant estimates (95% confidence intervals do not include zero).

| Directed edge | Boot.p | Opp.boot.p | Direct.p | Edge strength | Mean nonzero | SD nonzero | Lower 2.5% | Upper 97.5% |
|---|---|---|---|---|---|---|---|---|
| River → EggSize | 1 | 0 | 1 | 1 | **0.1951** | 0.0247 | 0.1467 | 0.2435 |
| River → FL | 1 | 0 | 1 | 1 | **-0.2371** | 0.0158 | -0.2680 | -0.2062 |
| FL → EggSize | 1 | 0 | 1 | 1 | **0.5377** | 0.1494 | 0.2450 | 0.8305 |
| FL → Fecund | 1 | 0 | 1 | 1 | **0.6204** | 0.0712 | 0.4809 | 0.7599 |
| JpChumN → RusChumN | 0.819 | 0.181 | 0.819 | 1 | **-0.7865** | 0.1285 | -1.0384 | -0.5346 |
| Fecund → EggSize | 0.642 | 0.358 | 0.642 | 1 | **-0.5490** | 0.0166 | -0.5815 | -0.5166 |
| NPGO3 → JpChumN | 0.984 | 0 | 1 | 0.984 | **0.3929** | 0.1554 | 0.0883 | 0.6975 |
| River → Fecund | 0.983 | 0 | 1 | 0.983 | **0.0794** | 0.0401 | 0.0007 | 0.1581 |
| RusChumN → FL | 0.971 | 0 | 1 | 0.971 | **-0.1962** | 0.0827 | -0.3583 | -0.0341 |
| NPGO3 → RusChumN | 0.905 | 0 | 1 | 0.905 | 0.1198 | 0.2554 | -0.3808 | 0.6204 |
| JpChumN → EggSize | 0.827 | 0 | 1 | 0.827 | **0.1141** | 0.0383 | 0.0389 | 0.1893 |
| JpChumN → FL | 0.766 | 0 | 1 | 0.766 | -0.1168 | 0.0904 | -0.2939 | 0.0603 |
| NPGO3 → FL | 0.690 | 0 | 1 | 0.690 | 0.0738 | 0.0644 | -0.0524 | 0.2001 |
| RusChumN → EggSize | 0.665 | 0 | 1 | 0.665 | **-0.0809** | 0.0399 | -0.1592 | -0.0026 |

Boot.p: probability that the directed edge was present in the bootstrap networks in the specified direction.
Opp.boot.p: probability that the reverse directed edge was present.
Direct.p: conditional probability of the specified direction given that either direction was present, calculated as Boot.p / (Boot.p + Opp.boot.p).

**Table S4.** Second-stage bootstrap estimates of standardized path coefficients of the 14 directed edges retained in the final Bayesian network and the eight mediated effects conditional on the final Bayesian network structure (Fig. 5). Mean and SD summarize the corresponding bootstrap distributions from 1,000 bootstrap replicates. For mediated pathways, the indirect effect was calculated as the product of the standardized path coefficients along the constituent edges. The 95% bootstrap confidence intervals were estimated from the 2.5th and 97.5th percentiles of the corresponding bootstrap distributions. Bold values indicate statistically significant estimates (95% confidence intervals do not include zero).

| Path way | Mean boot | SD boot | Lower 2.5% | Upper 97.5% |
|---|---|---|---|---|
| Directed pathway | | | | |
| River → EggSize | **0.2881** | 0.0292 | 0.2338 | 0.3489 |
| River → FL | **-0.3810** | 0.0271 | -0.4335 | -0.3292 |
| FL → EggSize | **0.6806** | 0.0719 | 0.5464 | 0.8245 |
| FL → Fecund | **0.4776** | 0.0628 | 0.3554 | 0.6001 |
| JpChumN → RusChumN | **-0.8089** | 0.1178 | -1.0434 | -0.5840 |
| Fecund → EggSize | **-0.6094** | 0.0728 | -0.7609 | -0.4610 |
| NPGO3 → JpChumN | **0.4401** | 0.1514 | 0.1041 | 0.6977 |
| River → Fecund | **-0.0724** | 0.0220 | -0.1186 | -0.0308 |
| RusChumN → FL | **-0.3172** | 0.1401 | -0.6239 | -0.0428 |
| NPGO3 → RusChumN | 0.1724 | 0.1771 | -0.1896 | 0.4904 |
| JpChumN → EggSize | **0.1714** | 0.0806 | 0.0133 | 0.3237 |
| JpChumN → FL | -0.1601 | 0.1429 | -0.4371 | 0.1412 |
| NPGO3 → FL | 0.1123 | 0.0919 | -0.0780 | 0.2824 |
| RusChumN → EggSize | -0.1215 | 0.0717 | -0.2747 | 0.0129 |
| Mediated pathway | | | | |
| River → FL × FL → EggSize | **-0.2589** | 0.0296 | -0.3270 | -0.2054 |
| River →Fecund × Fecund →EggSize | **0.0437** | 0.0131 | 0.0183 | 0.0711 |
| FL →Fecund × Fecund →EggSize | **-0.2934** | 0.0646 | -0.4344 | -0.1786 |
| RusChumN →FL × FL →EggSize | **-0.2173** | 0.1015 | -0.4334 | -0.0251 |
| JpChumN →RusChumN × RusChumN →FL | **0.2575** | 0.1261 | 0.0358 | 0.5558 |
| JpChumN →RusChumN × RusChumN → EggSize | 0.1014 | 0.0664 | -0.0098 | 0.2456 |
| NPGO3 →JpChumN × JpChumN →EggSize | 0.0729 | 0.0428 | -0.0034 | 0.1646 |
| NPGO3 →JpChumN × JpChumN →RusChumN | **-0.3664** | 0.1567 | -0.6944 | -0.0714 |

**Table S5.** Bootstrap estimates and 95% confidence intervals for standardized path coefficients of the 19 directed edges retained in the Bayesian network including Russian pink salmon abundance (RusPinkN), based on edge strength > 0.6 and direction probability > 0.5. Mean and SD summarize the bootstrap distribution of non-zero standardized coefficients from networks in which the corresponding directed edge was present. The 95% bootstrap confidence intervals were estimated using the normal approximation as mean ± 1.96 SD. Bold values indicate statistically significant estimates (95% confidence intervals do not include zero).

| Directed edge | Boot.p | Opp.boot.p | Direct.p | Edge strength | Mean nonzero | SD nonzero | Lower 2.5% | Upper 97.5% |
|---|---|---|---|---|---|---|---|---|
| River →EggSize | 1 | 0 | 1 | 1 | **0.1948** | 0.0244 | 0.1469 | 0.2427 |
| River →FL | 1 | 0 | 1 | 1 | **-0.2385** | 0.0152 | -0.2684 | -0.2087 |
| FL →EggSize | 1 | 0 | 1 | 1 | **0.5176** | 0.1522 | 0.2192 | 0.8160 |
| FL →Fecund | 1 | 0 | 1 | 1 | **0.6251** | 0.0748 | 0.4784 | 0.7718 |
| JpChumN →RusChumN | 0.871 | 0.129 | 0.871 | 1 | **-0.7829** | 0.1405 | -1.0584 | -0.5074 |
| Fecund →EggSize | 0.569 | 0.431 | 0.569 | 1 | **-0.5488** | 0.0165 | -0.5810 | -0.5165 |
| NPGO3 →JpChumN | 0.973 | 0 | 1 | 0.973 | **0.3913** | 0.1638 | 0.0702 | 0.7123 |
| River →Fecund | 0.971 | 0 | 1 | 0.971 | **0.0841** | 0.0423 | 0.0011 | 0.1670 |
| RusPinkN →FL | 0.963 | 0 | 1 | 0.963 | **-0.1606** | 0.0580 | -0.2744 | -0.0468 |
| RusChumN →RusPinkN | 0.540 | 0.420 | 0.563 | 0.960 | 0.4274 | 0.3371 | -0.2333 | 1.0881 |
| RusChumN →FL | 0.941 | 0 | 1 | 0.941 | **-0.1576** | 0.0742 | -0.3031 | -0.0120 |
| JpChumN →RusPinkN | 0.731 | 0.188 | 0.795 | 0.919 | -0.0693 | 0.3527 | -0.7607 | 0.6220 |
| NPGO3 →RusPinkN | 0.906 | 0 | 1 | 0.906 | -0.2998 | 0.2900 | -0.8682 | 0.2686 |
| NPGO3 →RusChumN | 0.902 | 0 | 1 | 0.902 | 0.1896 | 0.2384 | -0.2777 | 0.6569 |
| JpChumN →EggSize | 0.836 | 0 | 1 | 0.836 | **0.1110** | 0.0403 | 0.0319 | 0.1900 |
| RusPinkN →Fecund | 0.800 | 0 | 1 | 0.800 | **-0.0624** | 0.0259 | -0.1132 | -0.0116 |
| JpChumN →FL | 0.764 | 0 | 1 | 0.764 | -0.1346 | 0.0835 | -0.2983 | 0.0292 |
| RusChumN →EggSize | 0.736 | 0 | 1 | 0.736 | **-0.0868** | 0.0416 | -0.1682 | -0.0053 |
| NPGO3 →FL | 0.605 | 0 | 1 | 0.605 | 0.0284 | 0.0857 | -0.1397 | 0.1964 |

Boot.p: probability that the directed edge was present in the bootstrap networks in the specified direction.
Opp.boot.p: probability that the reverse directed edge was present.
Direct.p: conditional probability of the specified direction given that either direction was present, calculated as Boot.p / (Boot.p + Opp.boot.p).

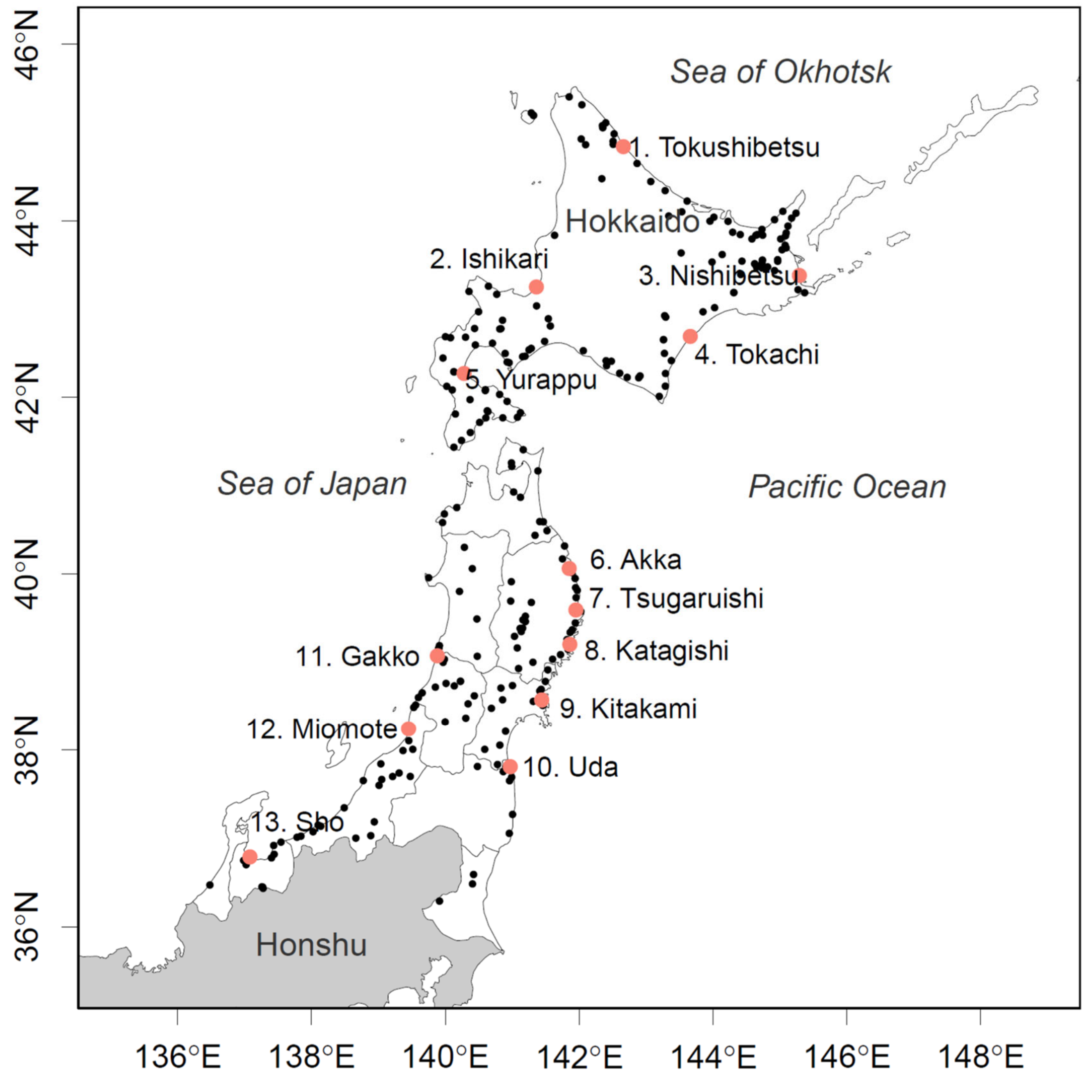


**Fig. S1.** Locations of 13 hatchery-enhanced rivers in Japan from which chum salmon were sampled. Red dots indicate the mouths of the sampled rivers, while the chum salmon return area is shown in white. Black dots indicate salmon hatcheries. Adapted from Fig. 2 in Kitada (2020).

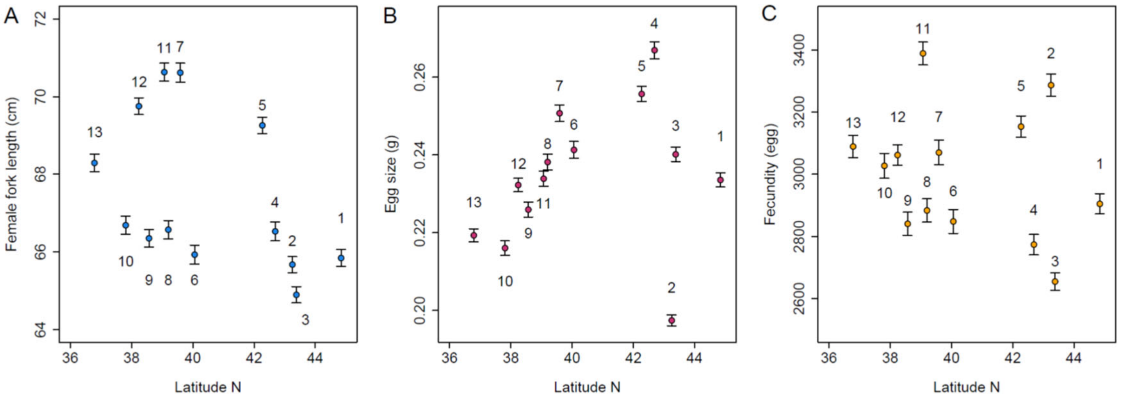


**Fig. S2.** Geographic distributions of mean female fork length (FL) (A), mean egg size (B), and fecundity (C) of age-4 hatchery-origin chum salmon sampled during 1999–2019 ($n$ = 12,428). Blue symbols in panels A represent river-specific mean FL, red symbols in panels B represent river-specific mean egg size, and orange symbols in panels C represent river-specific mean fecundity; error bars indicate 95% confidence intervals of the river-specific means calculated from all screened individual observations within each river during the study period. Numbers indicate the locations of the 13 sampling rivers (Fig. S1). Panels (A) and (B) were adapted from Figure 1 in Kitada (2026). The correlations between River and FL ($r = -0.46$, $t = -1.72$, df = 11, $p = 0.114$), River and fecundity ($r = -0.23$, $t = -0.80$, df = 11, $p = 0.440$), and River and egg size ($r = 0.23$, $t = 0.78$, df = 11, $p = 0.453$) were not statistically significant. Excluding three northernmost rivers—Tokushibetsu, Ishikari, and Nishibetsu (#1–3), egg size showed a strong geographic cline among the remaining rivers ($r = 0.93$, $t = 7.38$, df = 8, $p < 0.0001$).

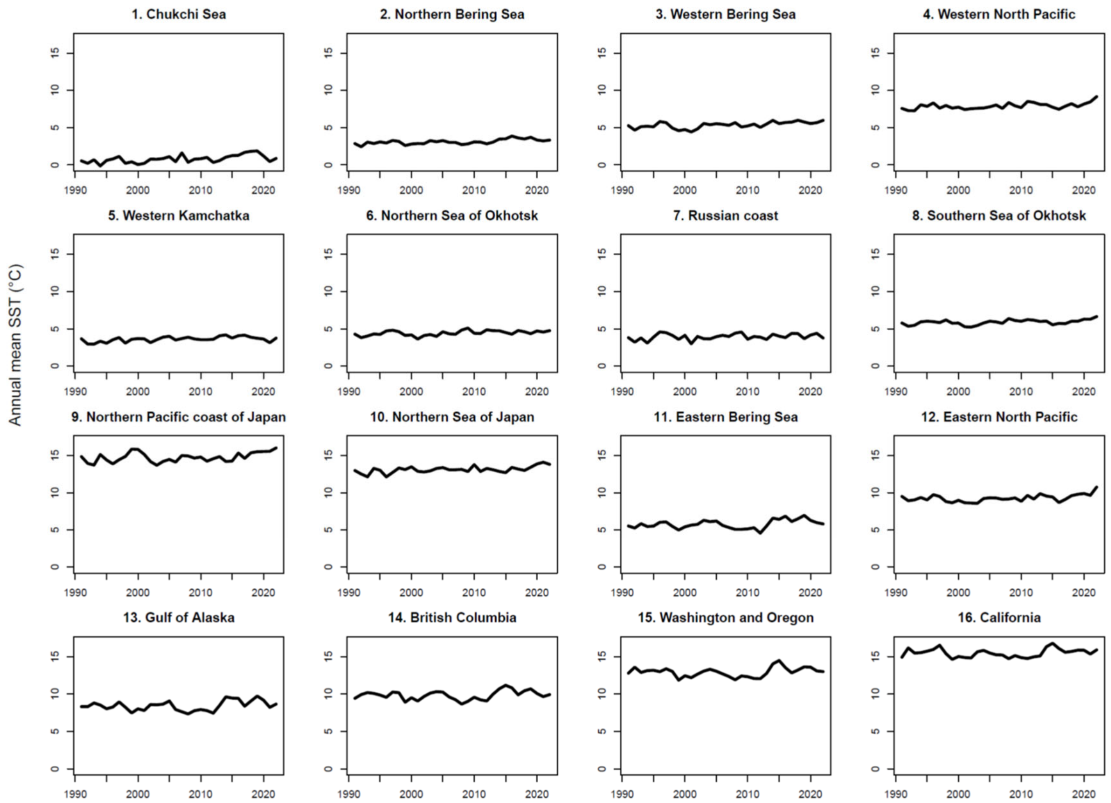


**Fig. S3.** Annual mean sea surface temperature (SST) across 16 areas within the chum salmon distribution range during 1991–2022 (see Fig. 2A).

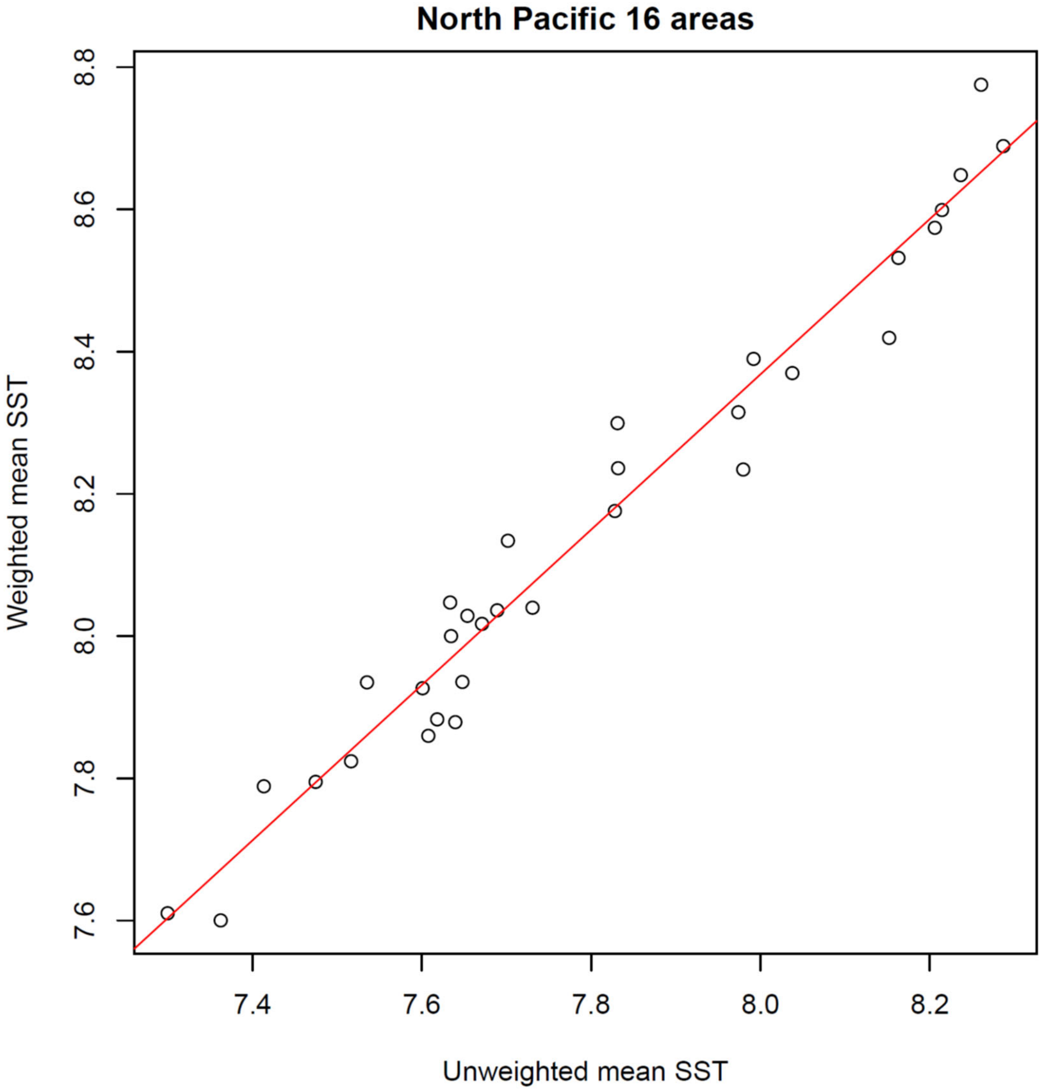


**Fig. S4.** Comparison of area-weighted and unweighted mean sea surface temperature (SST) indices for the 16 geographic areas shown in Fig. 2A. Each point represents an annual mean for 1991–2022. The red dashed line indicates the 1:1 relationship ($y = x$). The two indices were highly correlated ($r = 0.98$, $t = 30.38$, df = 30, $p < 0.0001$).

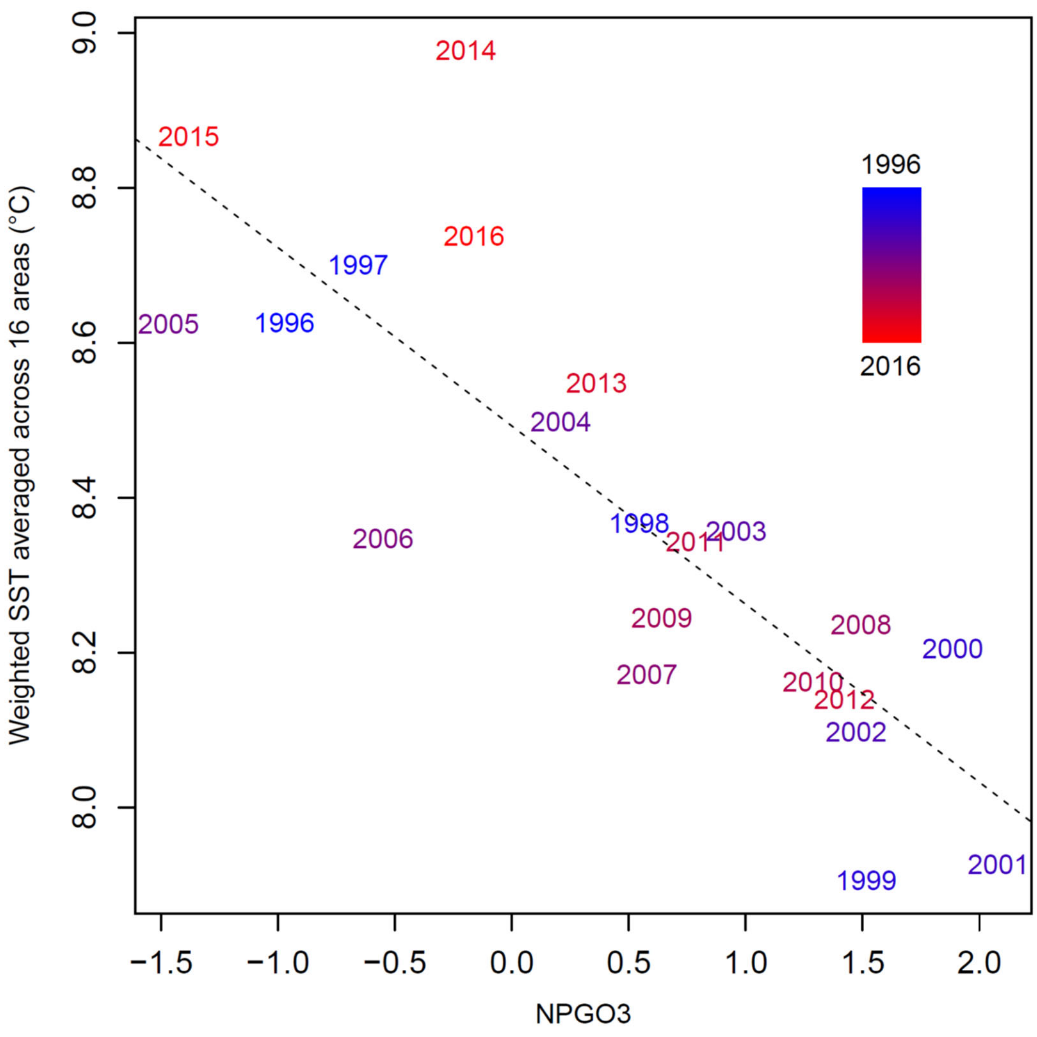


**Fig. S5.** Relationship between the North Pacific Gyre Oscillation (NPGO3) and annual weighted mean SST across the 16 areas in the release year of age-4 fish ($t$−3) (1996–2016) during the Bayesian network study period (1999–2019) (see Fig. 2B) ($r = -0.83$, $t = -6.42$, df = 19, $p < 0.0001$, adjusted $R^2 = 0.67$).

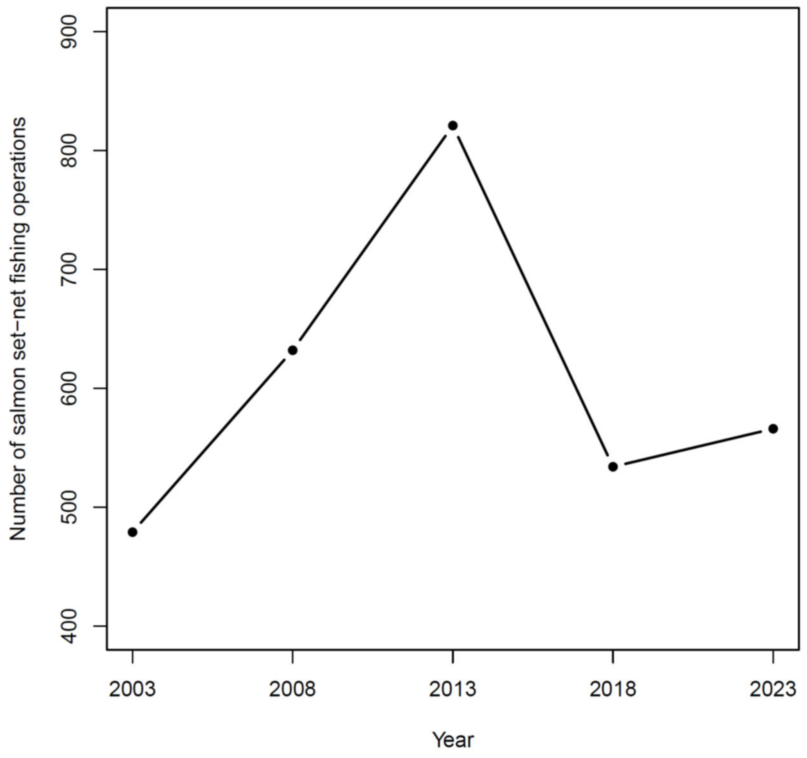


**Fig. S6.** Number of fishing operations engaged in chum salmon set-net fisheries in Japan from 2003 to 2023, based on the national fisheries census conducted at five-year intervals (MAFF 2026b). The mean number of fishing operations across the five census years was 606.

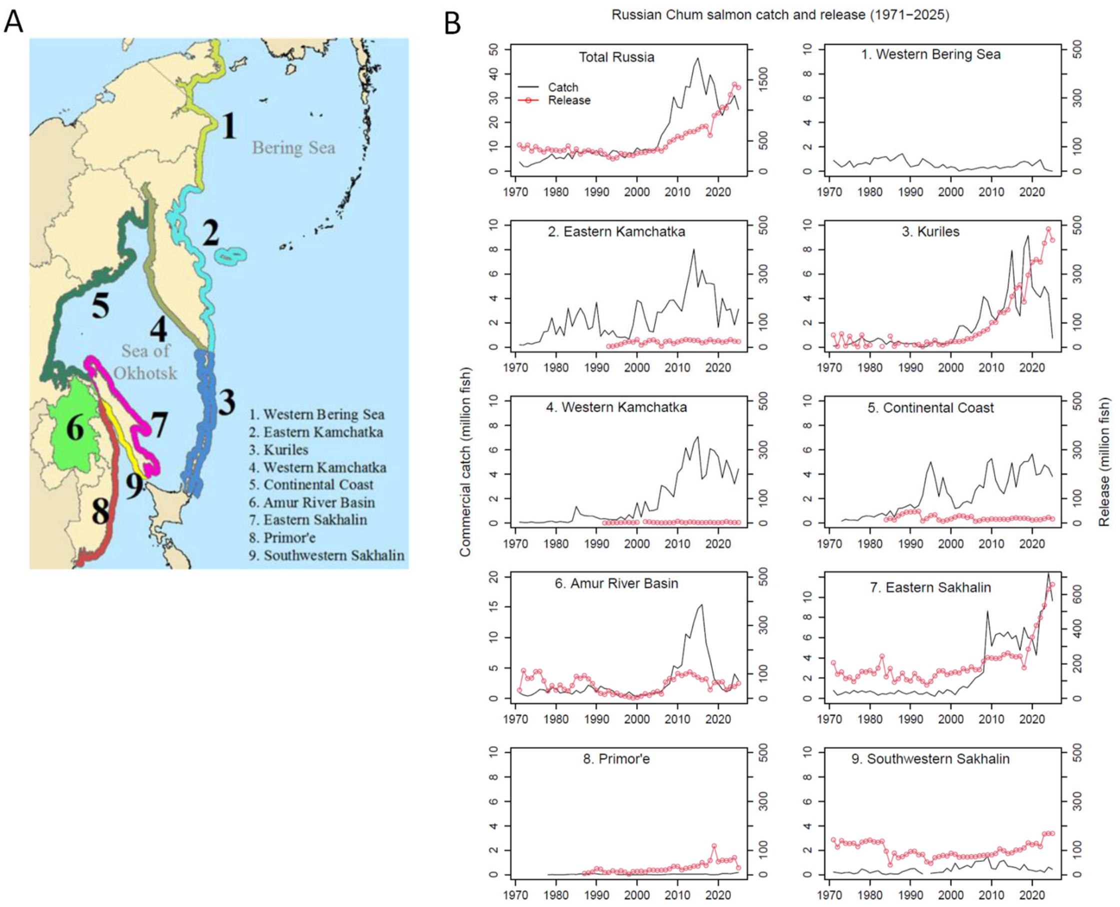


**Fig. S7.** Chum salmon commercial catch and hatchery release. (A) Russia reporting areas for salmon catch and hatchery releases (adapted from NPAFC 2026). (B) Changes in chum salmon commercial catch and hatchery release in the reporting areas (1971-2025).

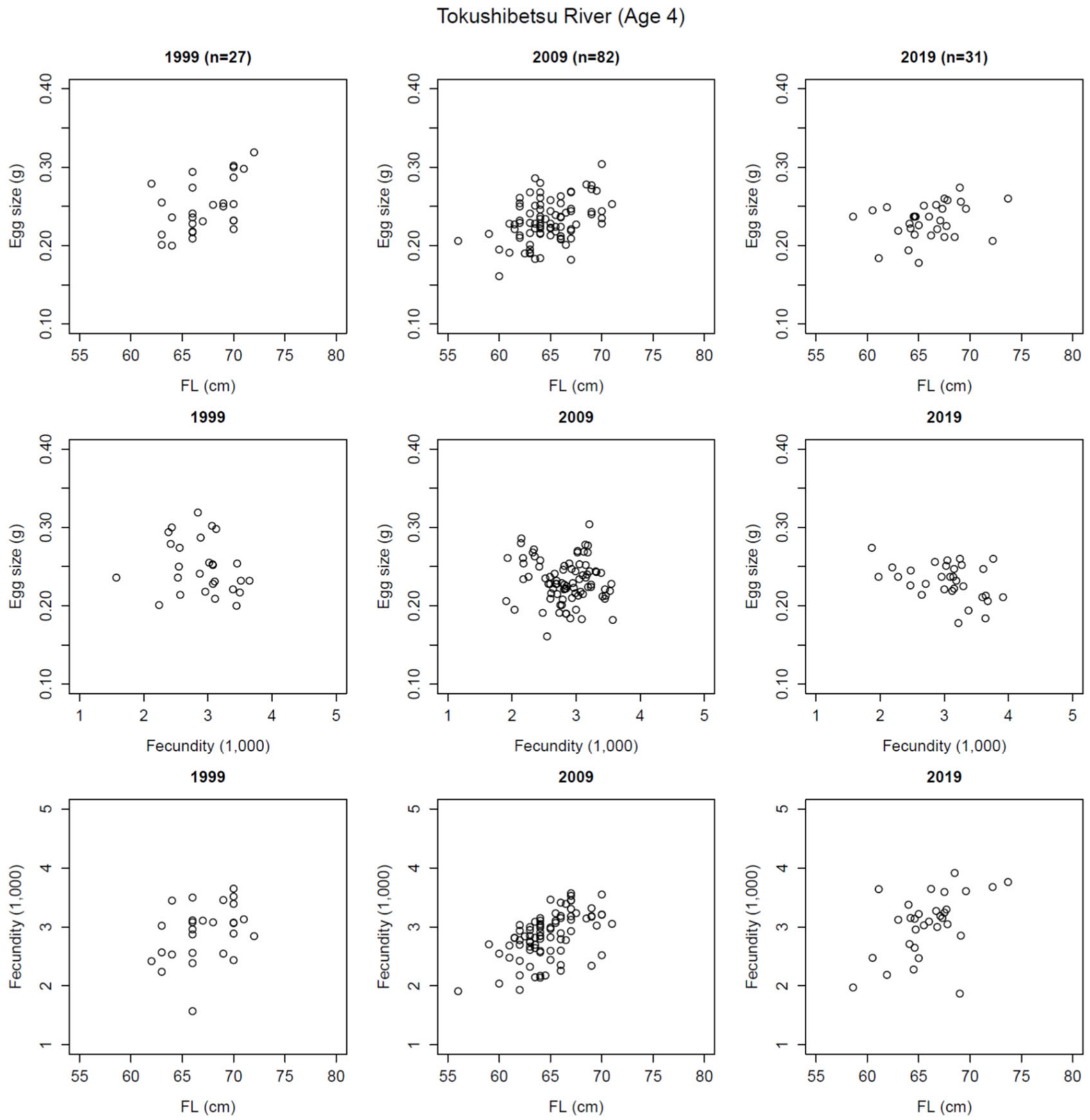


**Fig. S8.** Decadal changes in the relationships between fork length (FL) and reproductive traits of age-4 female chum salmon in the Tokushibetsu River, based on data before outlier removal ($n$ = 892).

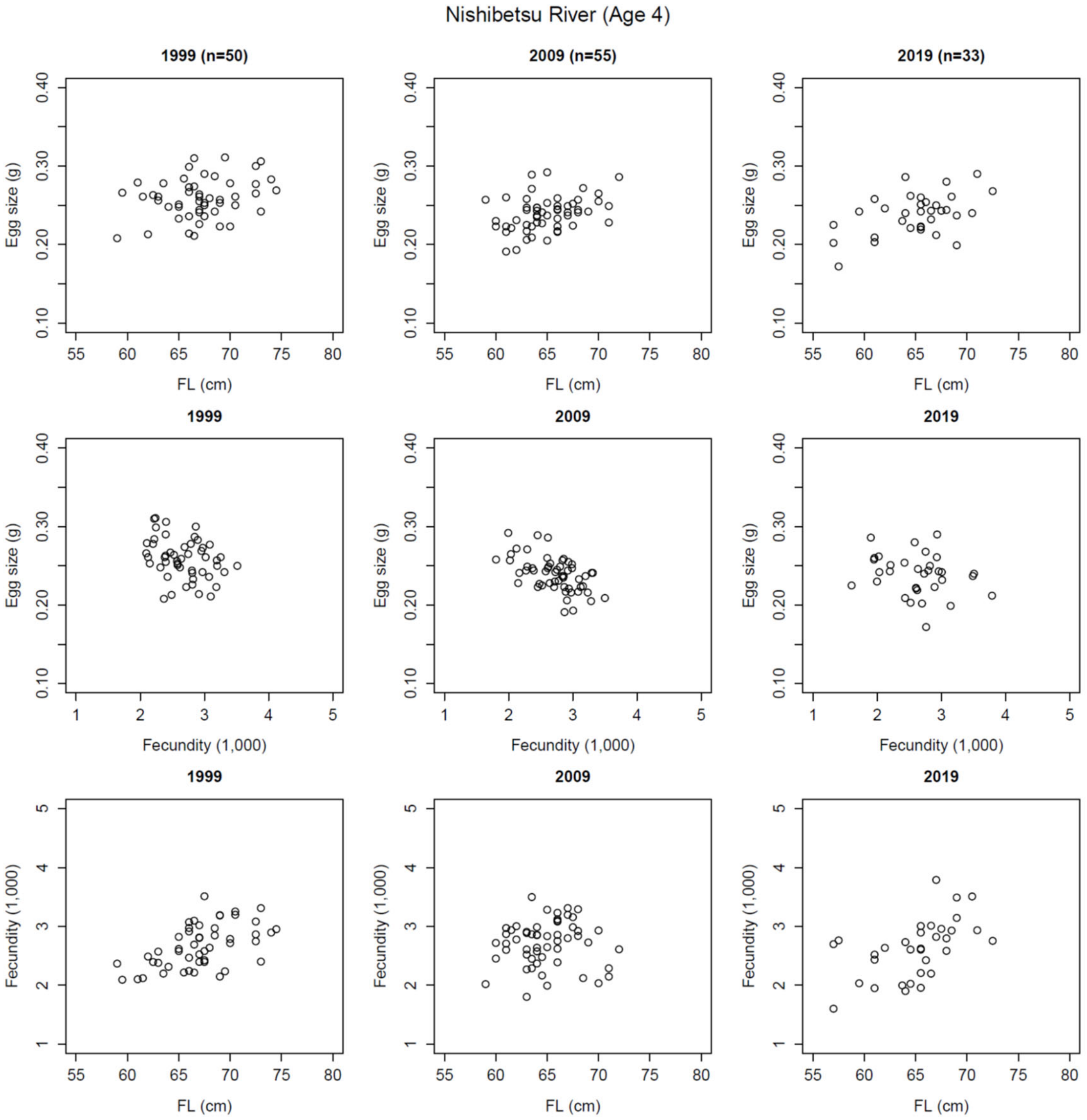


**Fig. S9.** Decadal changes in the relationships between fork length (FL) and reproductive traits of age-4 female chum salmon in the Nishibetsu River, based on data before outlier removal ($n$ = 1,071).

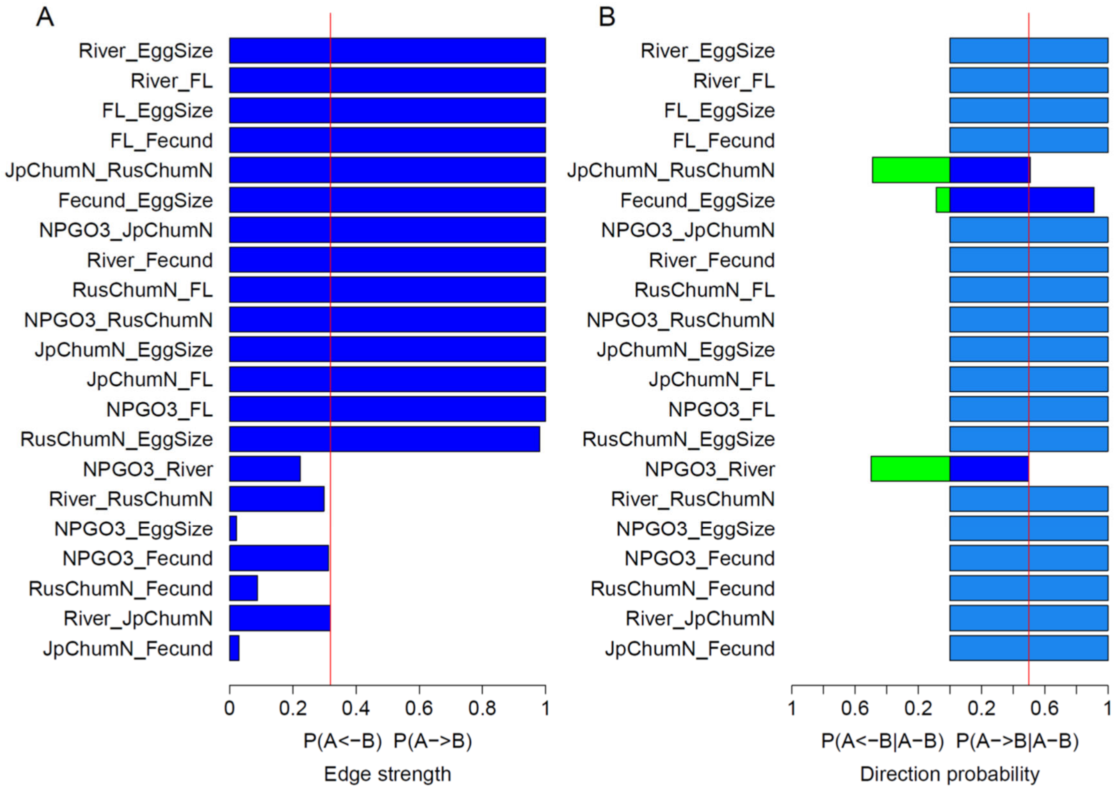


**Fig. S10.** Bootstrap support for Bayesian network edges estimated from 1,000 bootstrap replicates without accounting for temporal dependence and synchronous variation among rivers. (A) Edge strength. The vertical red line indicates the edge-strength threshold (0.314). The edge strengths showed a clear gap, with all edges above the gap; all retained edges had strengths ≥0.97, far exceeding the estimated inclusion threshold of 0.314. (B) Edge direction probability. Blue and green bars represent edge direction probabilities estimated without biological constraints, whereas light blue bars represent the corresponding estimates obtained under the biological constraints defined in Table S2.

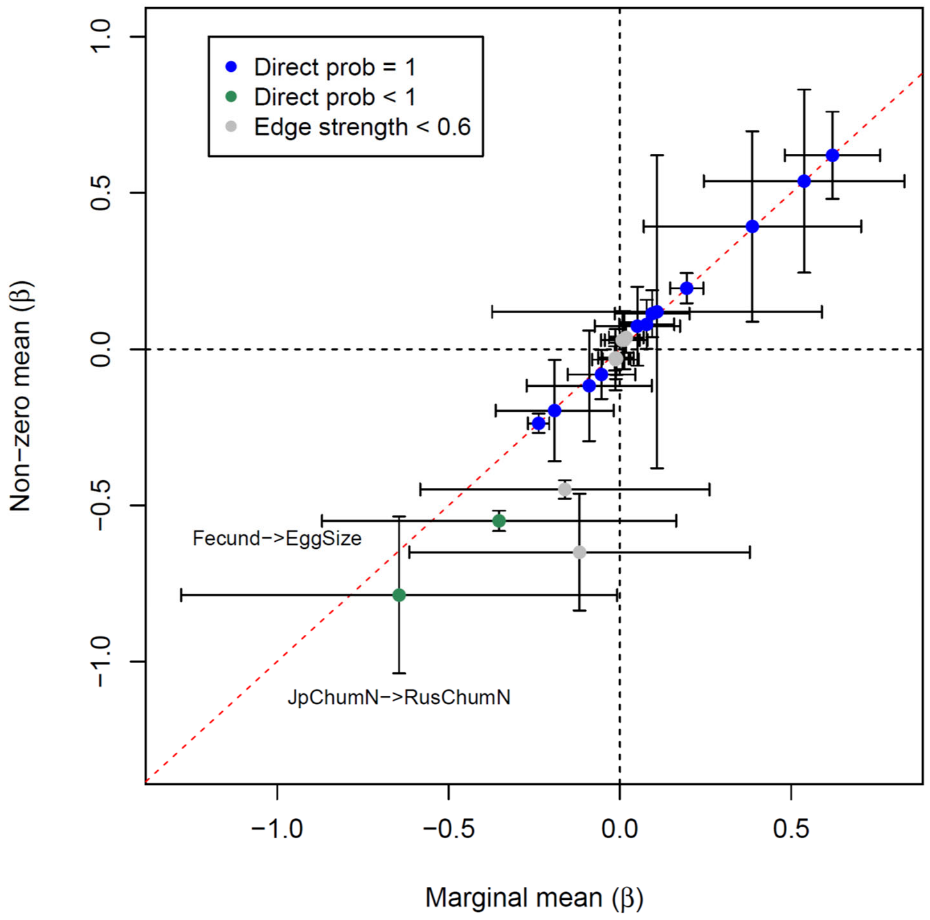


**Fig. S11.** Comparison of marginal and nonzero-mean bootstrap estimates of standardized path coefficients for the 23 directed edges shown in Fig. 4B. Points show the mean coefficient estimates, with error bars indicating the corresponding 95% confidence intervals. The nonzero-mean and marginal estimates were highly correlated ($r = 0.94$, $t = 12.15$, df = 21, $p < 0.0001$). Blue and green points indicate the directed edges retained in the final Bayesian network. Blue points indicate the 19 directed edges with a direction probability of 1.0, for which the two point estimates and their 95% confidence intervals were nearly identical. Green points indicate the two directed edges with lower direction probabilities, JpChumN → RusChumN (0.819) and Fecund → EggSize (0.642), for which the nonzero-mean estimates were smaller and had narrower 95% confidence intervals than the corresponding marginal estimates. Gray points indicate the two directed edges with direction probabilities below 0.5, RusChumN → JpChumN and EggSize → Fecund, which were excluded from the final network. The dashed red line represents the 1:1 relationship. The horizontal and vertical dashed lines indicate zero values for the directed effects and nonzero-mean path coefficients, respectively.

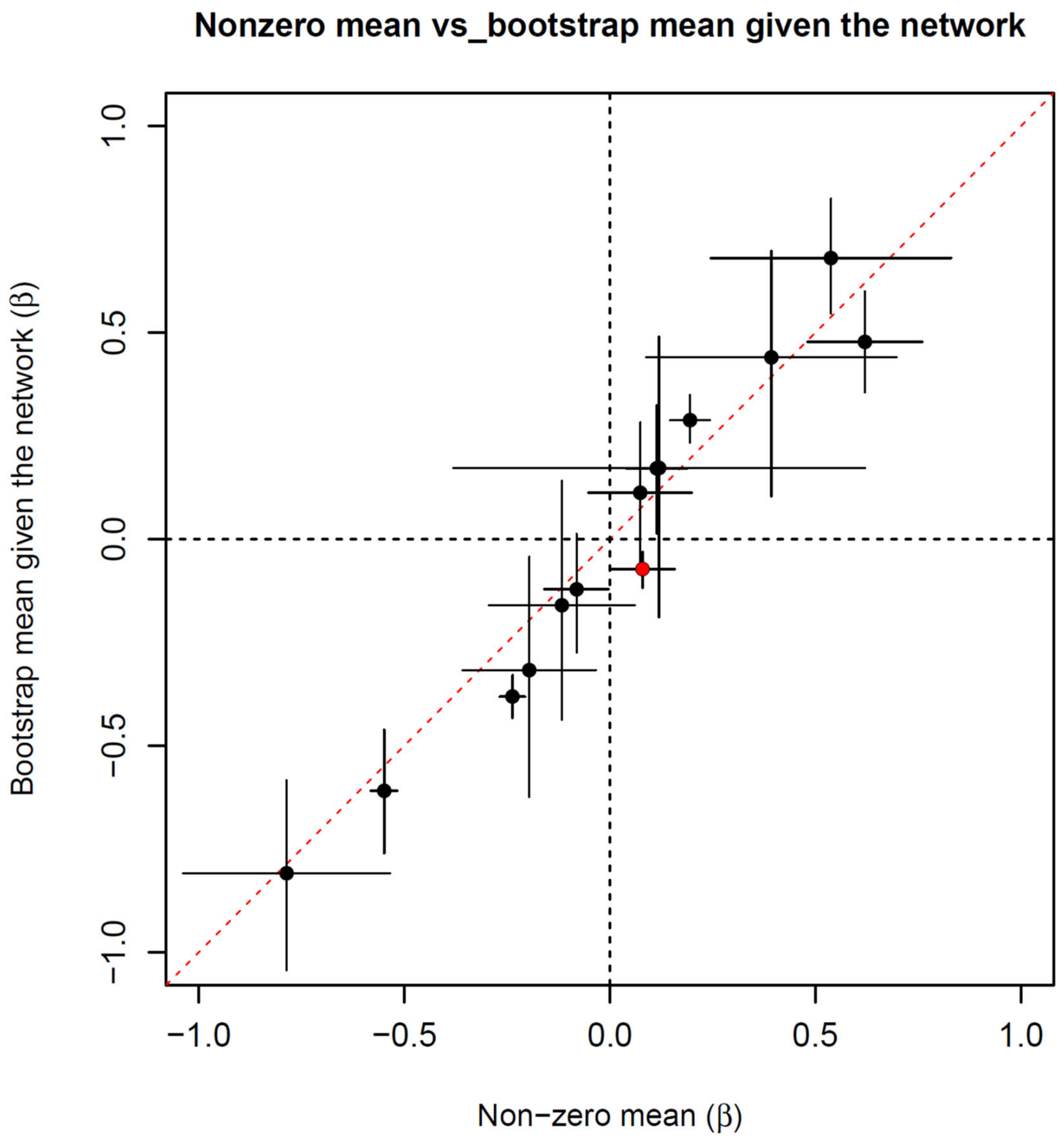


**Fig. S12.** Relationship between directed effects estimated from the second-stage bootstrap and corresponding nonzero-mean standardized path coefficients from the first-stage bootstrap. Points show the mean coefficient estimates of the 14 retained directed edges, with error bars indicating the corresponding 95% confidence intervals. The two estimates were highly correlated ($r$ = 0.98, $t$ = 15.83, df = 12, $p$ < 0.0001). The dashed red line represents the 1:1 relationship. The River → Fecund edge (red circle) was the only edge for which the signs of the two estimates differed. The horizontal and vertical dashed lines indicate zero values for the directed effects and nonzero-mean path coefficients, respectively.

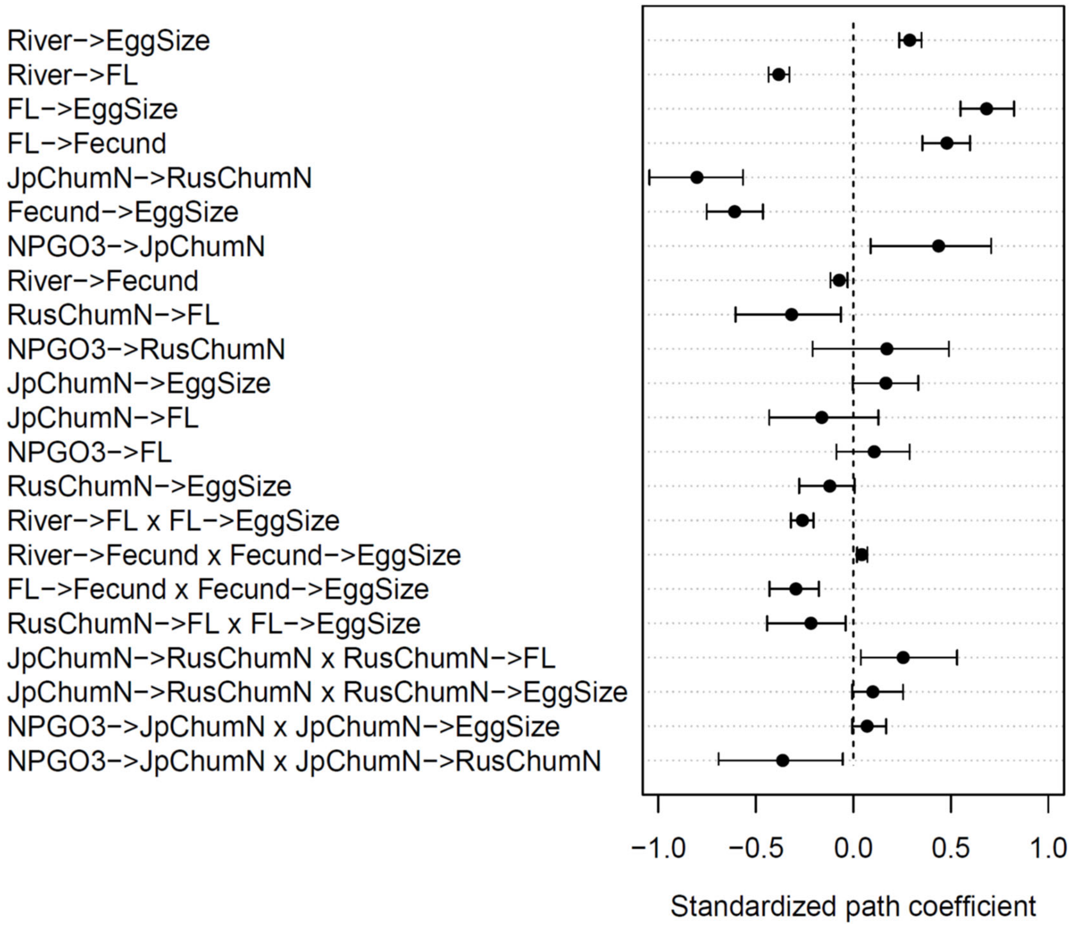


**Fig. S13.** Bootstrap estimates of standardized path coefficients (β) and mediated effects for the 14 directional edges and 8 selected mediated pathways in the final Bayesian network (see Fig. 5). Dots indicate bootstrap means, and horizontal bars represent 95% bootstrap confidence intervals based on the 2.5th and 97.5th percentiles of the corresponding bootstrap distributions across 1,000 second-stage bootstrap replicates (Table S4). For mediated pathways, indirect effects were calculated as the products of the standardized path coefficients along the constituent edges. Estimates whose confidence intervals exclude zero were considered statistically supported.

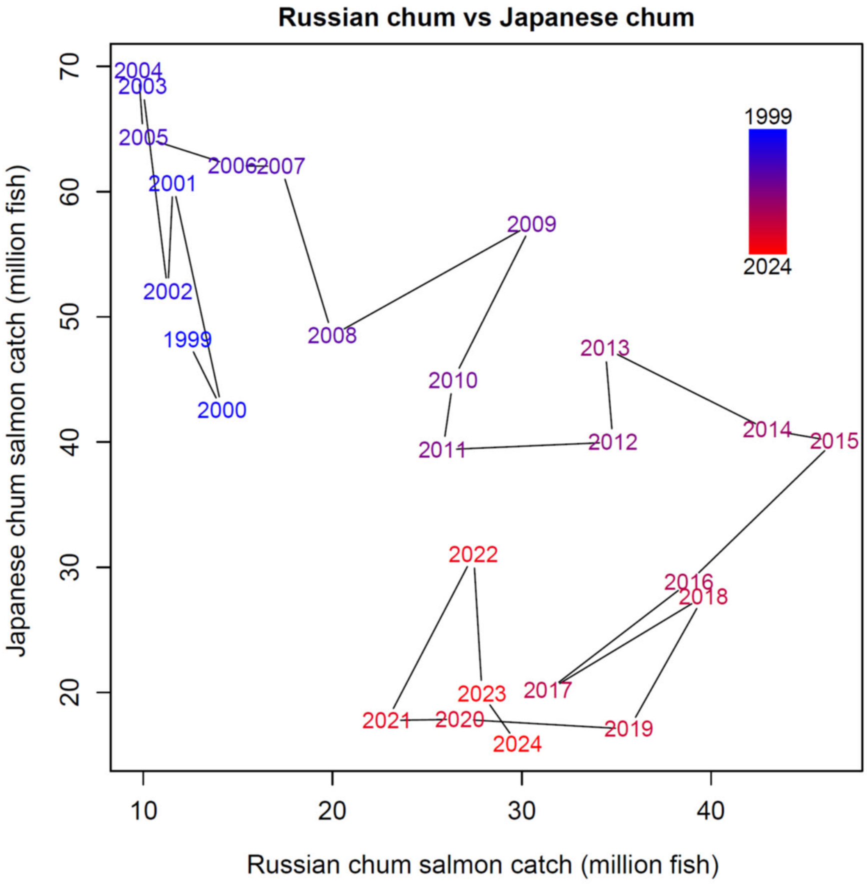


**Fig. S14.** Temporal relationship between Russian and Japanese chum salmon catches (1999−2024). A significant negative correlation was observed ($r = -0.61$, $t = -3.78$, df = 24, $p = 0.0009$); however, this relationship reversed following a pronounced decline in Russian chum abundance.

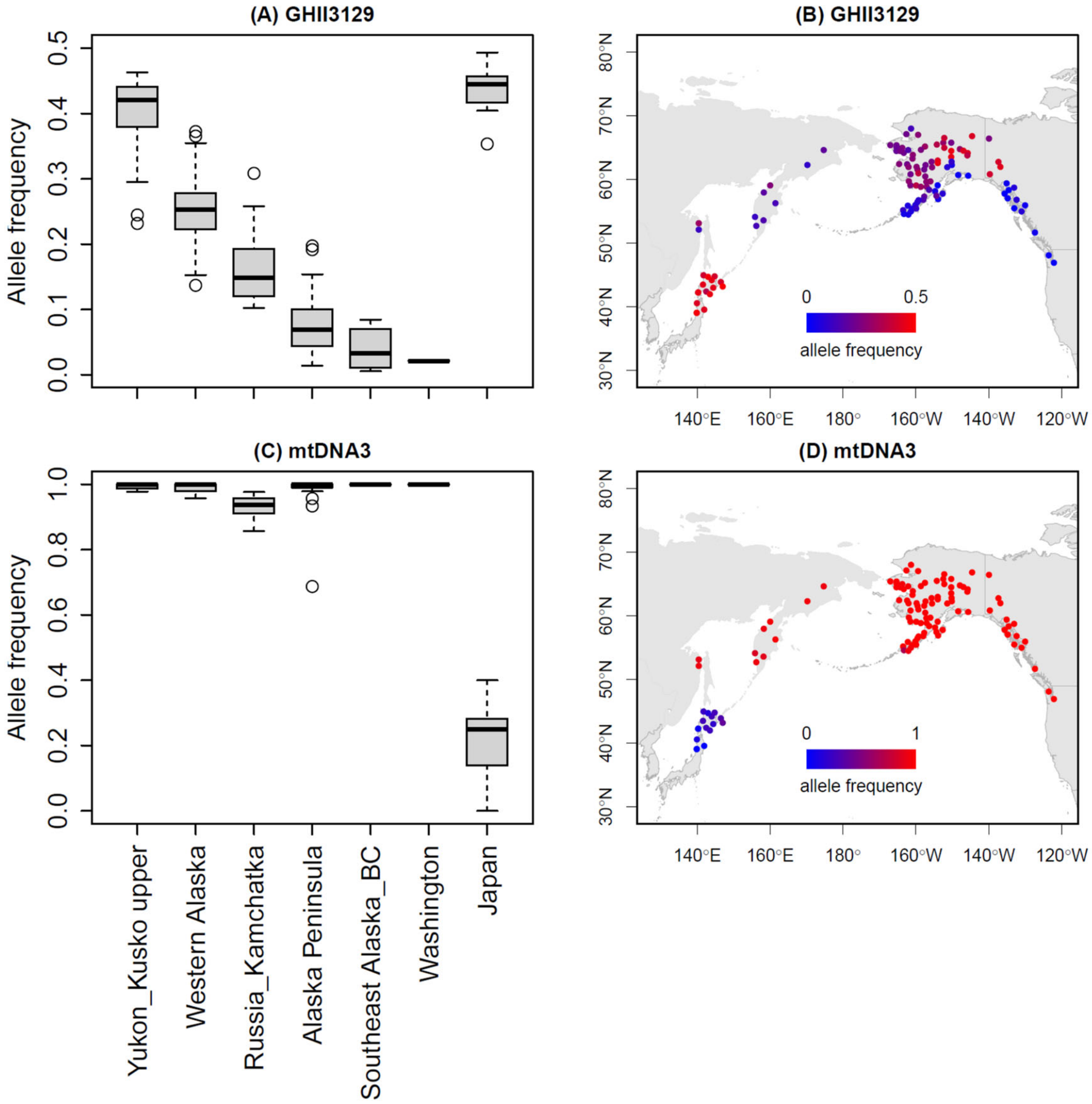


**Fig. S15.** Geographic distributions of allele frequencies at two genetic loci in Japanese and North Pacific chum salmon populations. (A, B) Growth hormone locus Oke_GHII-3129; (C, D) combined mitochondrial DNA locus mtDNA3. (A, C) Boxplots of allele frequencies by sampling region, arranged from north (left) to south (right). (B, D) Geographic distributions of allele frequencies at 114 sampling locations, with point colors proportional to allele frequency. (A, B) were newly prepared using data from Kitada and Kishino (2021); (C, D) were adapted from Fig. 4 of Kitada and Kishino (2021). At Oke_GHII-3129, allele frequency generally declined from approximately 0.42 in the northern populations of the Yukon and upper Kuskokwim rivers to below 0.05 in Washington, whereas Japan showed a relatively high allele frequency of approximately 0.45. In contrast, mtDNA3 allele frequency remained high (approximately 0.9–1.0) from Alaska to Washington but was substantially lower in Japan (approximately 0.2).

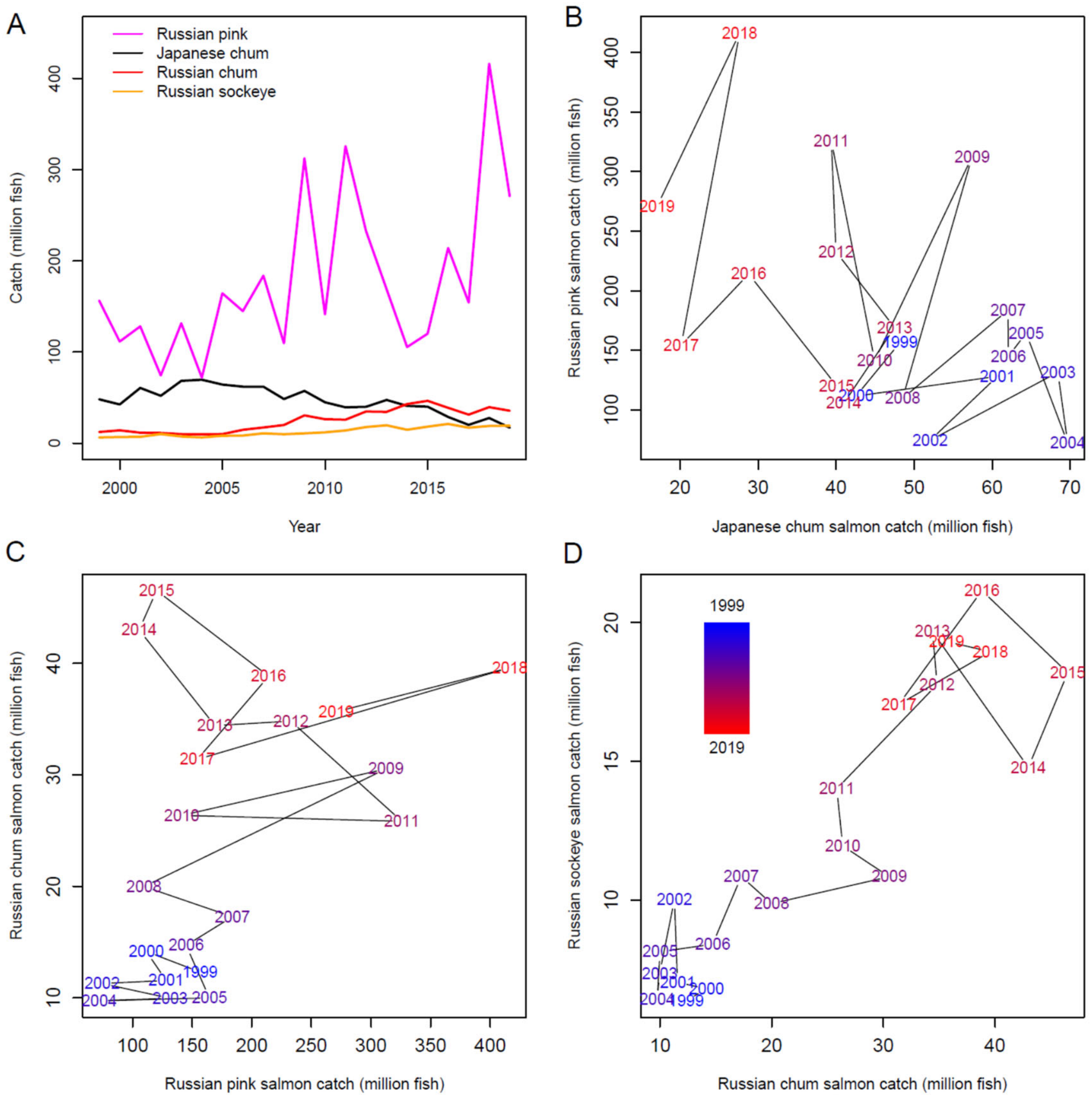


**Fig. S16.** Temporal trends and pairwise relationships among salmon catches in Japan and Russia during 1999–2019. (A) Annual catches of Japanese chum salmon, Russian chum salmon, Russian pink salmon, and Russian sockeye salmon. (B) Relationship between Japanese chum salmon and Russian pink salmon catches. (C) Relationship between Russian pink salmon and Russian chum salmon catches. (D) Relationship between Russian chum salmon and Russian sockeye salmon catches. Japanese chum salmon and Russian pink salmon catches were moderately negatively correlated ($r = -0.43$, $p = 0.049$), whereas Russian chum salmon and Russian pink salmon catches were moderately positively correlated ($r = 0.44$, $p = 0.044$). Russian chum salmon and Russian sockeye salmon catches were strongly positively correlated ($r = 0.90$, $p < 0.001$).

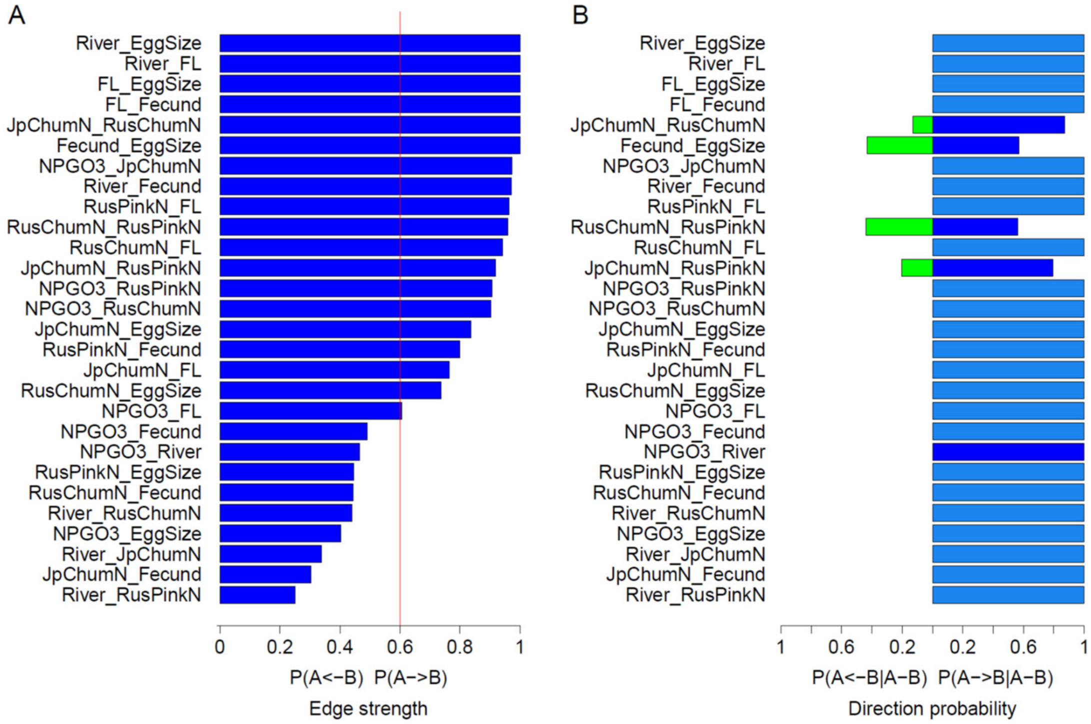


**Fig. S17.** Bootstrap support for Bayesian network edges estimated from 1,000 block-bootstrap replicates with Russian pink salmon abundance (RusPinkN) included as an additional node. Years were resampled while retaining all individuals ($n$ = 12,428) within each year, thereby accounting for temporal dependence and synchronous variation among rivers. (A) Edge strength. The vertical red line indicates the edge-strength threshold (0.6). Edges with edge strength >0.6 and direction probability >0.5 were retained for construction of the Bayesian network shown in Fig. 8. (B) Edge direction probability. Blue and green bars represent edge direction probabilities estimated without biological constraints, whereas light blue bars represent the corresponding estimates obtained under the biological constraints defined in Table S2. Green bars indicate edges with direction probabilities <0.5 and were therefore excluded from the network.

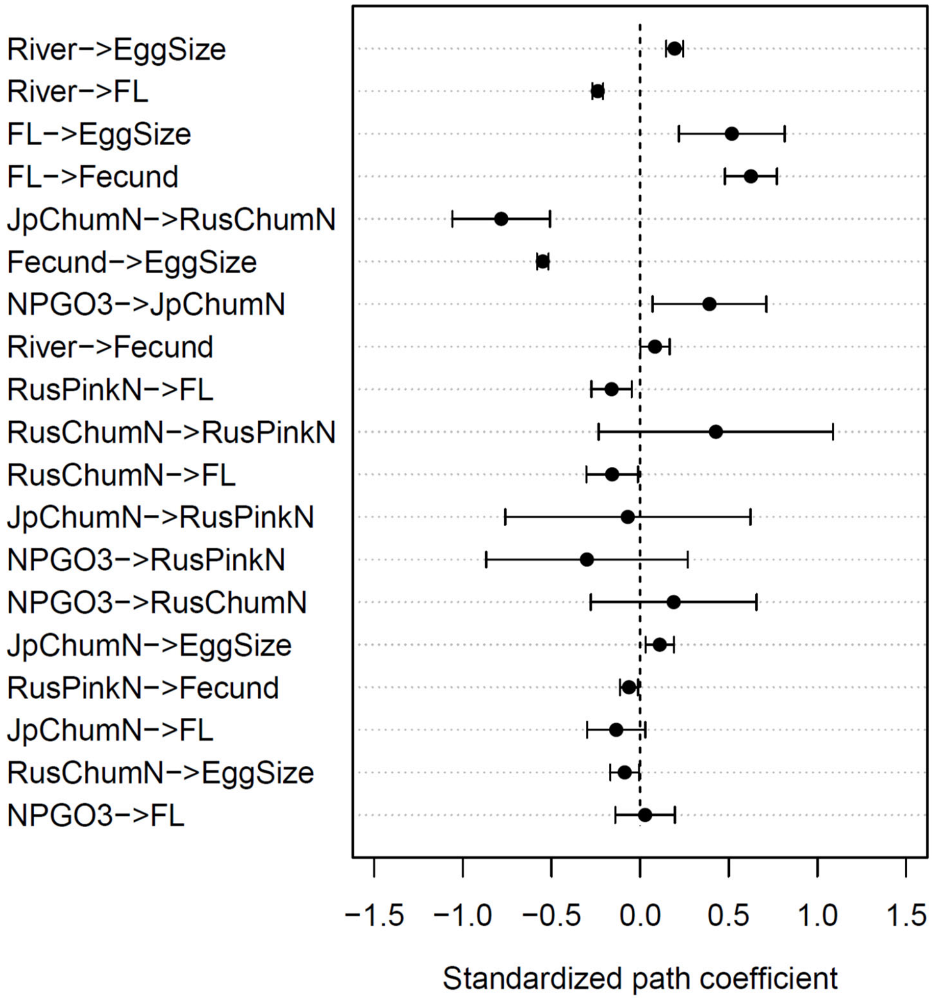


**Fig. S18.** Bootstrap-averaged nonzero-mean standardized path coefficients (β) for the directional edges in the Bayesian network including Russian pink salmon abundance (RusPinkN; see Fig. 8). Dots indicate bootstrap means, and horizontal bars represent 95% bootstrap confidence intervals based on the normal approximation (mean ± 1.96 SD) across 1,000 block bootstrap replicates in which years (1999–2019) were resampled while retaining all individuals within each year ($n$ = 12,428). Edges whose confidence intervals exclude zero were considered statistically supported.

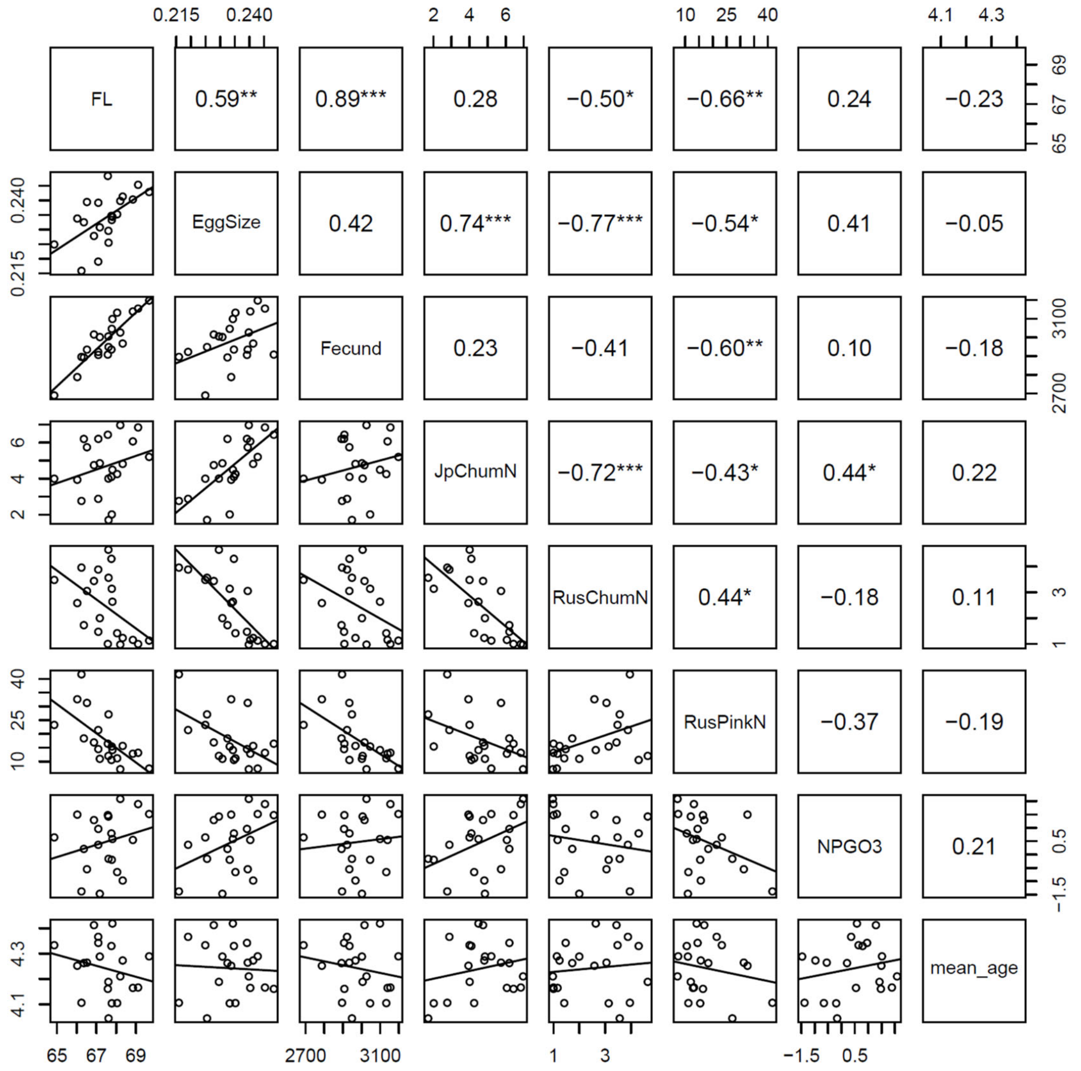


**Fig. S19.** Pairwise relationships among the six annual variables included in the final Bayesian network (excluding River), Russian pink salmon abundance (RusPinkN), and mean age at return during 1999–2019. The lower triangle shows scatter plots with fitted linear regression lines, and the upper triangle shows Pearson correlation coefficients. Asterisks indicate statistical significance ($p < 0.05$, $*p < 0.01$, $**p < 0.001$).

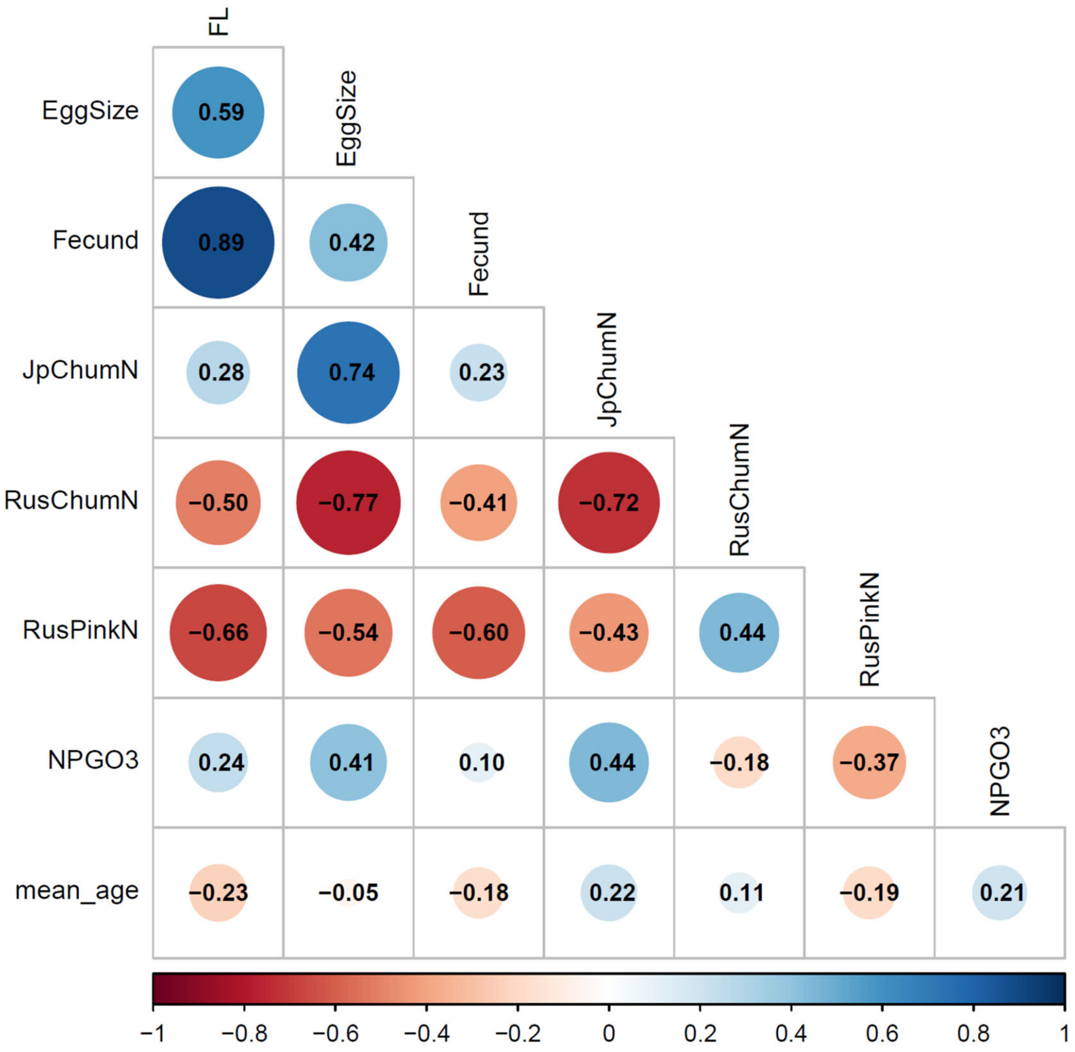


**Fig. S20.** Visual representation of pairwise Pearson correlation coefficients among the same eight variables shown in Fig. S19 during 1999–2019. Circle size indicates the magnitude of the Pearson correlation coefficient, and circle color indicates its direction, with blue representing positive and red representing negative correlations.

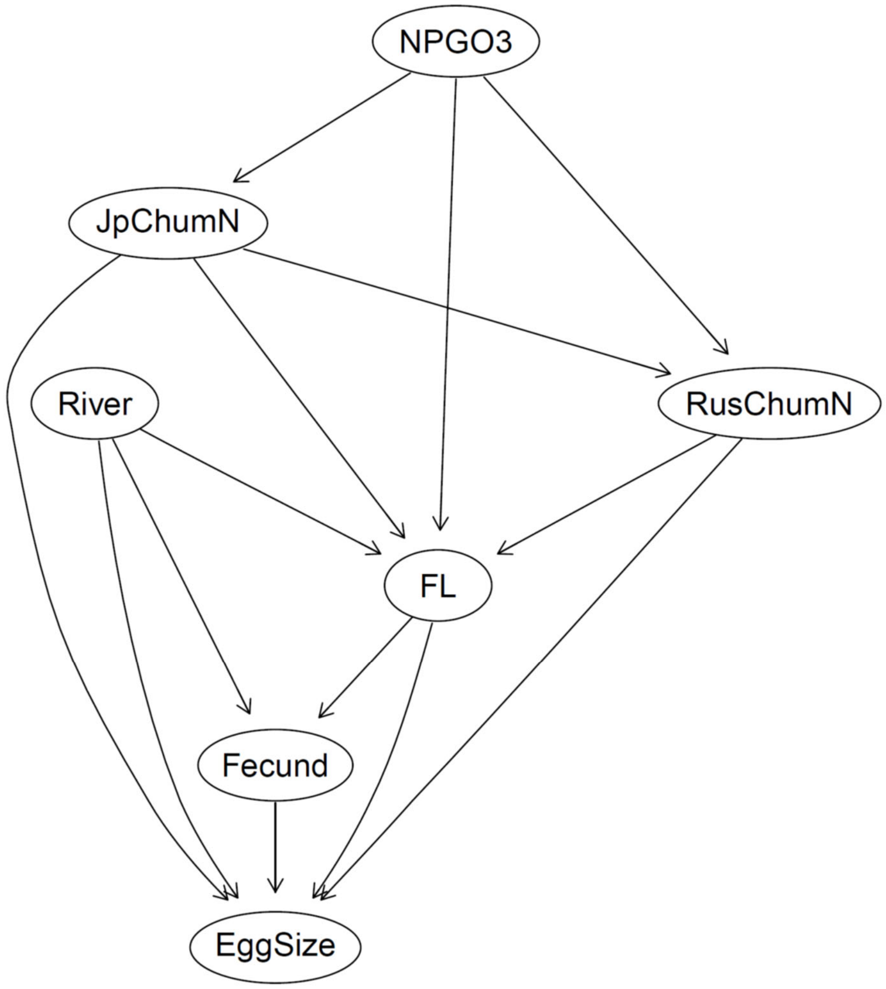


**Fig. S21.** Averaged Bayesian network without accounting for temporal dependence among rivers obtained using the `averaged.network()` function in the `bnlearn` package. Edge strengths and direction probabilities were estimated using the `boot.strength()` function based on 1,000 bootstrap replicates performed without accounting for temporal dependence among observations across rivers. The inclusion threshold was estimated using the `inclusion.threshold()` function (0.31). The averaged network was constructed from edges with strengths exceeding the threshold (> 0.31) and direction probabilities greater than 0.5. A direction probability > 0.5 indicates that the inferred direction was recovered more frequently than the opposite direction across bootstrap replicates. Despite not accounting for temporal dependence, the resulting network was identical to the bootstrap-supported final Bayesian network shown in Fig. 5, consisting of seven nodes and 14 directed edges.

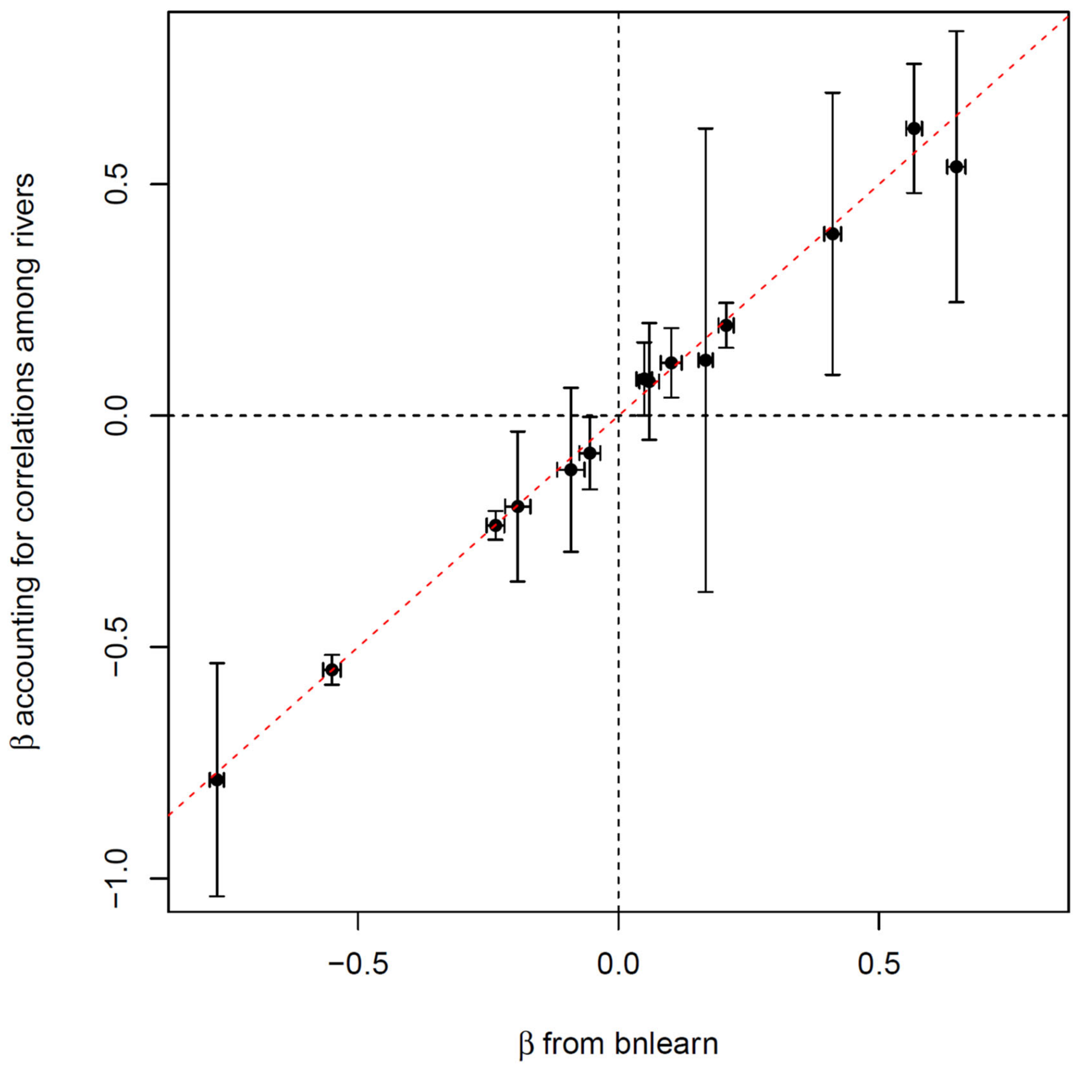


**Fig. S22.** Comparison of standardized path coefficients (β) for the 14 directed edges estimated from the bootstrap-based Bayesian network accounting for river-level correlation (Fig. 5; y-axis) and the averaged Bayesian network ignoring river-level correlation (Fig. S20; x-axis). Error bars represent standard errors. The point estimates are highly consistent between the two approaches ($r = 0.995$, $t = 35.37$, df = 12, $p < 0.0001$), whereas the averaged Bayesian network yields substantially smaller standard errors, suggesting that uncertainty may be underestimated when river-level correlation is ignored. The red line indicates the 1:1 relationship (y = x).

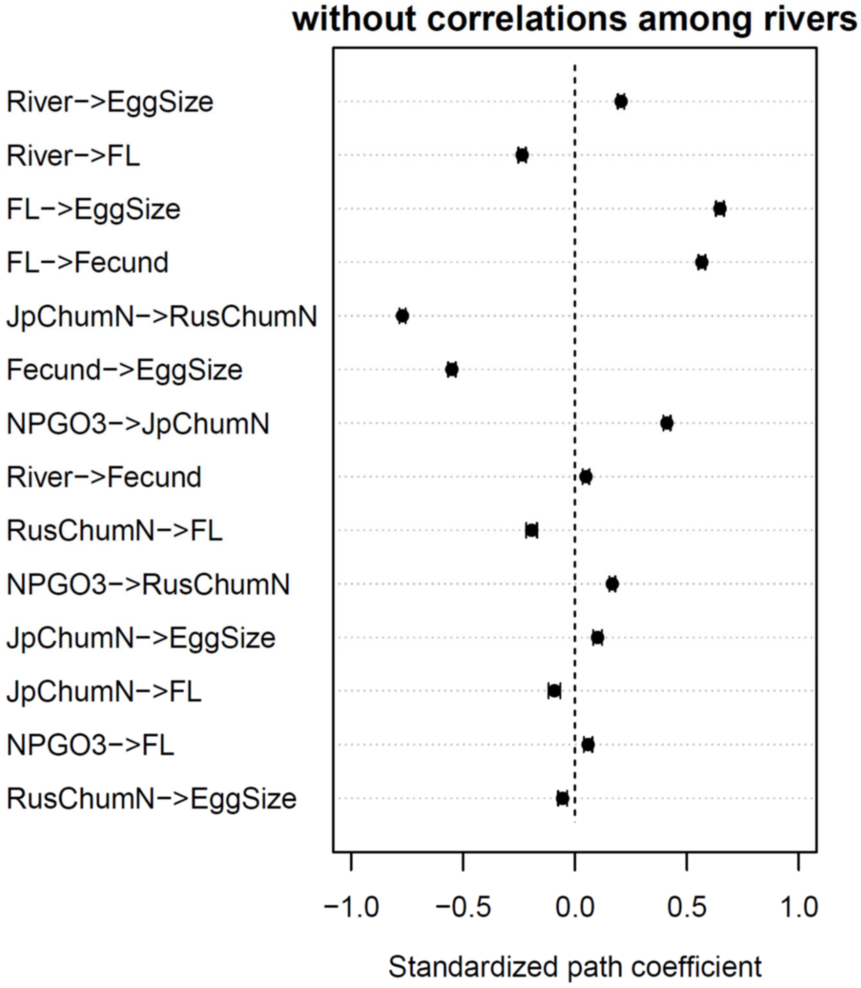


**Fig. S23.** Standardized path coefficients (β) for the 14 directed edges in the averaged Bayesian network without accounting for temporal dependence among rivers (Fig. S20). Dots indicate point estimates, and horizontal bars represent 95% confidence intervals (point estimate ± 1.96 SE). All confidence intervals are narrow and exclude zero.
